# Comparative and population genomics analyses of the regulatory Transcription Factor (TF)-DNA interaction

**DOI:** 10.1101/2024.11.05.622057

**Authors:** Manas Joshi, Pablo Duchen, Adamandia Kapopoulou, Stefan Laurent

**Affiliations:** Max-Planck-Institut für Pflanzenzüchtungsforschung, Germany; Institute of Organismic and Molecular Evolution, Johannes Gutenberg University Mainz, Germany; Institute of Ecology and Evolution, University of Bern, Switzerland; Swiss Institute of Bioinformatics, Switzerland

## Abstract

Natural selection heavily influences the evolutionary trajectories of species by impacting their genotype to phenotype transitions. On the molecular level, these genotype to phenotype transitions are shaped by the regulatory sequences (RSs). In this study, we employed a combination of population and comparative genomics to investigate how natural selection affects specific RS classes involved in the regulatory transcription factor (TF)-DNA interactions. These interactions consist of two motifs, namely: TF-binding domains (TF-BDs) and TF-binding sites (TF-BSs). Using publicly available annotation data, we constructed the species-specific lists of the TF-BD and TF-BS regions. We also constructed a non-synonymous equivalent class for the TF-BS regions for making comparative analyses across the coding and non-coding regions. On applying some of the commonly used summary statistics, we found a consistent signal of purifying selection acting on TF-BDs, consistent with their functional importance. Interestingly, we also observed that non-coding TF-BS regions showed similar levels of constraint to that of coding regions for populations with large N_e_. Concerning positive selection, we showed that the regulatory motifs of large N_e_ species are, overall, subject to an increased intensity of positive selection. We found an overall positive correlation between the intensity of purifying and positive selection and the species-specific Ne, which is expected under drift-selection equilibrium. Finally, we contrasted the signal of positive selection in *H. sapiens* and *D. melanogaster* (outcrossing species) with *A. thaliana* (selfing species) and noted that selfing plays a key role in influencing the action of positive selection.

## Introduction

The study of natural selection and how it impacts the genome has been at the forefront of evolutionary biology research for several decades (Ellegren, 2008; Vitti et al., 2013). Several studies have highlighted that selection directly affects given phenotypes, which, in turn, will contribute to the genetic changes associated with those phenotypes. However, different parts of the genome could be subjected to varying intensities of natural selection. For instance, functional classes like coding regions have been shown to be differentially affected by selection compared to non-coding regions (Haddrill et al., 2008; Naidoo et al., 2018; Torgerson et al., 2009). The intensity of selection can also vary within the functional coding classes, such as selection acting on synonymous versus non-synonymous sites (Parsch et al., 2010).

Transcription factor (TF)-DNA interactions constitute an important class of regulatory interactions that play a vital role in controlling the context-dependent expression of genes, thereby, impacting the genotype to phenotype transitions. These interactions act as the initial step in the transcriptional process of genes. Given their central role in controlling the expression patterns of the effector genes, these interactions, directly and indirectly, serve various functions ranging from cell differentiation and development (Lee & Young, 2013) to controlling multiple biological pathways (Desvergne et al., 2006). These regulatory interactions consist of two interacting motifs: TF-binding domains (TF-BDs) and TF-binding sites (TF-BSs). Here, TF-BDs are stretches of nucleotides within the protein-coding regions of TFs that code for the DNA-binding domains. On the other hand, TF-BSs are stretches of non-coding DNA that are usually located upstream of the effector gene to which the TFs bind via their TF-BD. These interacting motifs are stretches of regulatory sequences (RSs) which play an important role in gene regulation. Hence, variants occurring in these RSs could potentially alter the expression patterns of multiple effector genes, thereby influencing the genotype to phenotype transitions. Therefore, natural selection would be expected to act on these elements with an increased intensity compared to the genomic background.

Several studies have highlighted cases of TF-BS regions evolving under the pressure of natural selection when compared to the genomic background (reviewed in Joshi et al., 2021). Specifically, TF-BSs have been shown to have a reduced nucleotide diversity as compared to neutral classes, suggesting that these regions are under an increased intensity of purifying selection (Connelly et al., 2013; Mu et al., 2011; Vernot et al., 2012). Interestingly, Vernot et al., 2012, showed that binding sites for TFs enriched for cell differentiation and development processes showed particularly lower diversity, suggesting that purifying selection acts with varying intensity on different classes of TF-BSs. In addition to the action of purifying selection, studies have also highlighted that these regions are under the influence of positive selection (Arbiza et al., 2013; He et al., 2011). He et al., 2011 also attributed the species-specific gains and losses of TF-BS regions to positive selection. In summary, the evolution of the different classes of TF-BS regions has been documented to be driven by both positive and purifying selection. On the other hand, TF-BDs have been shown to be under increased phylogenetic conservation (Romani & Moreno, 2021). However, to the best of our knowledge, no previous studies have focused on elucidating the action of natural selection on TF-BDs through a molecular population genetics framework.

In this study, we set out to identify the impact of natural selection acting on both TF-BDs and TF-BSs through a comparative and population genetics framework. To this end, this study spanned three species, namely – *Homo sapiens*, *Arabidopsis thaliana* and *Drosophila melanogaster*, and six populations belonging to them (two populations per species). Importantly, the three species have varying magnitudes of effective population sizes (N_e_, *H. sapiens* < *A. thaliana* < *D. melanogaster*). This enabled us to perceive the observed selection signals through different drift-selection equilibria. Additionally, by adopting a combination of comparative and population genomics frameworks, we were able to study the action of selection on RSs over the two distinct evolutionary timescales. To make comparable analyses between the coding (TF-BDs and control regions) and non-coding (TF-BSs) regions, we constructed a “non-synonymous” equivalent class in the non-coding region based on their potential to disrupt the binding action of the TF-BDs and TF-BSs (a similar approach has been proposed in He et al., 2011). In TF-BDs, non-synonymous mutations are critical as they could disrupt multiple downstream regulatory interactions (Chesmore et al., 2016a). On the other hand, mutations occurring in the TF-BSs have also been annotated to have functional consequences (He et al., 2011; Mu et al., 2011). However, no previous study has performed a joint analysis to elucidate the impact of natural selection on the function-altering mutations occurring within both TF-BDs and TF-BSs. This is the first study that analyzed regulatory evolution using comparative and population genomic variation data spanning three species and their respective populations using annotated regulatory elements within both coding and non-coding regions of the genome. Our analyses noted a signal of high constraint acting on the TF-BDs compared to the TF-BSs and the control regions. We also noted that the TF-BS regions are overall under relaxed constraint when compared to the coding regions, which is in concordance with previous reports (Naidoo et al., 2018; Torgerson et al., 2009). However, in the case of large N_e_ species, we interestingly find that the TF-BS regions are under comparable levels of constraint as compared to certain coding classes. In terms of positive selection, we note that the overall signals of the RSs are under an increased intensity of positive selection compared to the control classes. Additionally, the three species involved in this study employ different mating strategies. Specifically, *H. sapiens* and *D. melanogaster* are outcrossing species, while *A. thaliana* is a selfing species. In selfing species, recombination rates are comparatively lower; hence, natural selection would be expected to act with lower efficiency. We note this influence of differences in selfing strategies in the positive selection analyses and further, confirm our findings through simulations using different selfing regimes.

## Materials and methods

### Divergence and Polymorphism analysis

For all the three species included in this study, *Homo sapiens*, *Arabidopsis thaliana*, and *Drosophila melanogaster*, we chose the following outgroup species: *Pan troglodytes*, *Arabidopsis lyrata*, and *Drosophila simulans*, respectively (Table 1). To generate alignments of TFs, we performed a gene-based orthology search using the reciprocal-blast approach. Specifically, every representative transcript in the ingroup species was aligned to the entire transcriptome from the outgroup species using *blastn* (Altschul et al., 1990) to obtain an outgroup transcript with the maximum identity score (minimum identity threshold – 60%). This outgroup transcript was then aligned back to the transcriptome of the ingroup species to investigate if the initial ingroup transcript had the maximum identity score. Then, the ascertained orthologous pair of transcript sequences from the ingroup and outgroup sequences were re-aligned with *MUSCLE* (Edgar, 2004). We also performed polymorphism-based analyses across six different populations from the three species. The species-specific sources of the polymorphism data are listed in (Table 1). Polymorphic sites within each population were polarized with the respective outgroup.

**Table 1.**
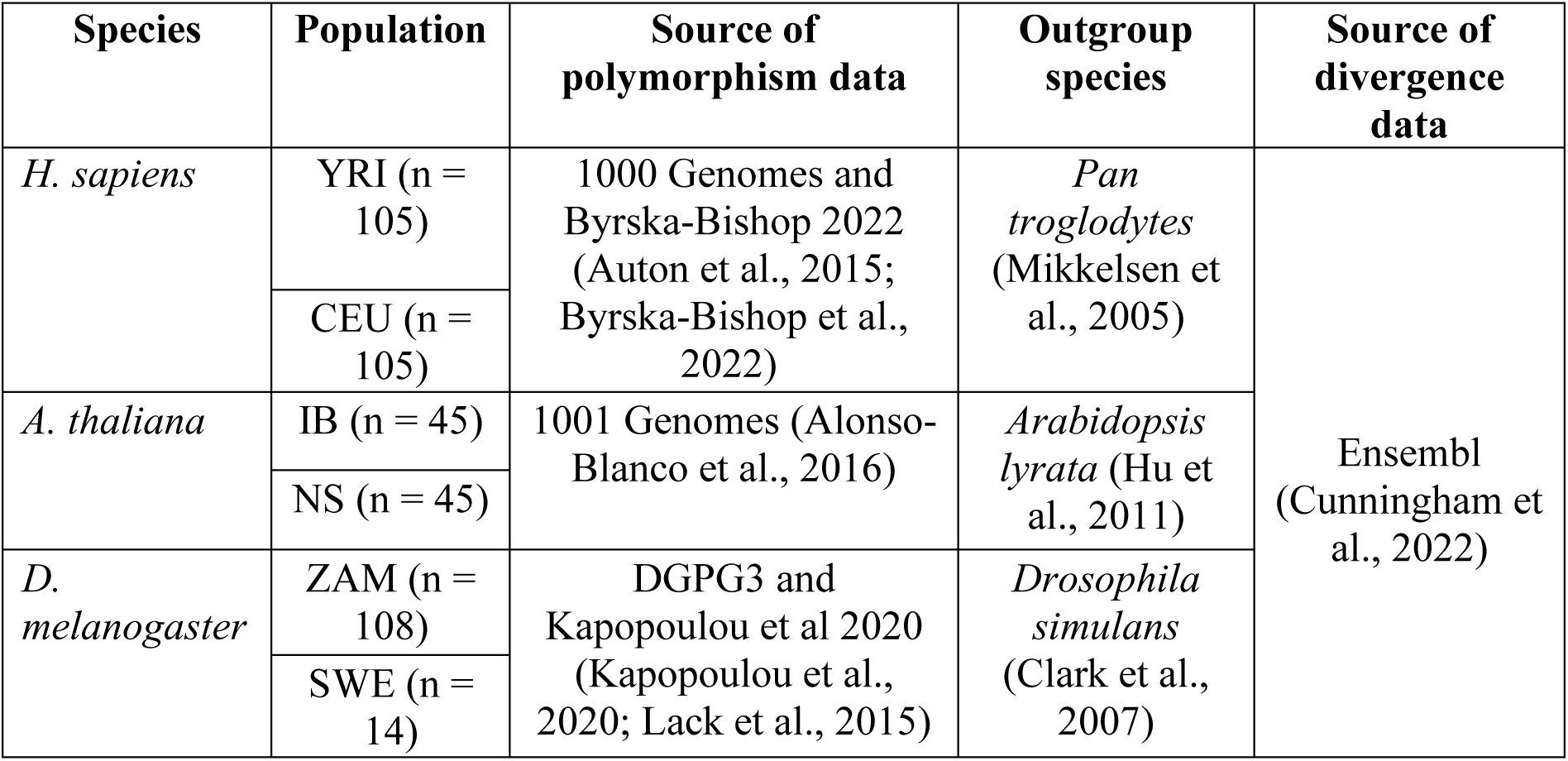
Overview of the polymorphism and divergence data used in this study. Population codes: YRI – Yoruba, CEU – Utah residents with Central European Ancestry, IB – Iberia, NS – North Sweden, ZAM – Zambia and SWE – Sweden. The population-specific sample sizes are mentioned in brackets.

### TF annotation and genomic regions included in this study

We constructed a species-specific list of TFs and the regulatory TF-BDs incorporated within them through annotations retrieved from UniProt (Bateman et al., 2023). To do this, we first extracted all the species-specific manually annotated (*Swissport identifiers*) protein entries and their respective annotated domains listed on UniProt. We further used domain-specific annotations to retain domains that were annotated with certain gene ontology (GO) terms indicating a regulatory role. The regulatory TF-BDs were identified based on the GO terms – “*Transcriptional regulation*”, “*Transcriptional activity*” and “*DNA binding*” (Supplementary Figure S1). Proteins containing at least one domain annotated with the keywords of interest were included in the species-specific TF list. This list of TFs and their corresponding domains was then used in further TF-BD analyses. We then generated a catalogue of the species-specific TF family distribution (by domains) included in this study (Supplementary Table ST1, ST2, ST3).

Overall, to contrast the signals of selection on the regulatory domains, we constructed two control regions. First, using the domain-specific annotation data per TF, we extracted the functionally unannotated regions within the TF-coding genes. In this study, these regions are referred to as non-BD (non-binding domains). Given the genomic proximity of the non-BD and TF-BD regions, those two regions would experience similar recombination rates, mutation rates, and GC content. We also used the entire protein coding gene set per species (which includes the TFs), referred to as All genes, as an additional control region. The All genes class aids in getting an overview of the intensity of selection acting on the coding regions of species, and contrasting it to their regulatory elements.

The choice of representative transcripts per gene varied based on the category of genes under investigation. For the TF-centered analyses (BD and non-BD regions), we identified a transcript that had a corresponding manually curated protein identifier in the UniProt database and chose this as the representative transcript. This curated protein entry in UniProt also contained information on the functional domain annotations, thereby enabling us to assign the domain annotation information to every gene. For the All genes analyses, the approach for choosing transcript varied depending on the species. Specifically, for *H. sapiens*, we chose MANE (Matched Annotation from NCBI and EMBL-EBI, Morales et al., 2022) transcripts as the representative transcripts. In the case of *A. thaliana* and *D. melanogaster*, we chose Ensembl canonical transcripts as the representative transcripts (Table 2).

**Table 2.** Overview of the species-specific number of genes included in the All genes class and the analysis-dependent choice of representative transcript. The choice of transcript for the BD and non-BD analysis was exclusively based on the available manually annotated Uniprot (*Swissprot*) identifiers. However, in the case of All genes analysis, this choice was species-dependent. The MANE transcript was chosen as the representative transcript per gene for the *H. sapiens* analysis, whereas for *A. thaliana* and *D. melanogaster* this choice was based on the canonical Ensembl transcript.

| Species | No. of All genes | Choice of transcript (BD and non-BD) | Choice of transcript (All genes) |
| --- | --- | --- | --- |
| *H. sapiens* | 17751 | Uniprot identifier | MANE |
| *A. thaliana* | 27419 | Uniprot identifier | Canonical |
| *D. melanogaster* | 17679 | Uniprot identifier | Canonical |

### Constraint ratios

We defined constraint ratios as the proportion of non-synonymous variants (*test* class variants) to the proportion of synonymous variants (*neutral* class variants). Here, the synonymous variants are used as a control class to counter various confounding factors like change in demography, bottleneck effect, etc. The within-species polymorphism constraint ratios were denoted by π_n_/π_s_ and the between-species divergence constraint ratios were denoted by K_n_/K_s_. Additionally, the proportions of non-synonymous (π_n_ and K_n_) and synonymous (π_s_ and K_s_) variants were normalized with their respective Nei-Gojobori distances (Nei & Gojobori, 1986). To isolate the effect of negative selection on the different functional classes and species we discarded admixed individuals. Additionally, to get an estimate of variance around the constraint ratios, we also performed 500 non-parametric resampling cycles to get the confidence intervals (CIs) around the estimates.

### Access to clinically annotated variants

To calculate the proportions of annotated deleterious variants for TF-BD and non-BD, we employed the ClinVar dataset (Landrum et al., 2018). In a nutshell, the ClinVar database catalogues human genomic variants that are annotated to have clinical significance. The variants were extracted through the ClinVar data repository (date of accession – 2021-10-16). These variants were first filtered to retain only SNPs located within the coding regions of the TFs contained in our dataset. These variants were further filtered to retain those having a clinical significance i.e. benign (annotated as – “benign” and “likely benign”) and pathogenic (annotated as – “pathogenic” and “likely pathogenic”). For every genomic region, the proportion of pathogenic variants was calculated by taking the ratio of the number of pathogenic variants to the total number of reported clinical variants (pathogenic and benign).

### Identifying species-specific TF-BS regions

We used the ReMap2022 dataset (Hammal et al., 2022) to identify potential TF-BS regions within the genomes of the three species. Specifically, ReMap2022 catalogues species-specific regions which are binding hotspots for multiple transcriptional regulators (TRs). These regions are identified within the database as cis-regulatory modules (CRMs). We used these CRMs as the starting points to identify potential TF-BS regions. We filtered these CRMs to keep those present exclusively in the 2kb vicinity of the coding regions and having an annotation score of more than 30 (Supplementary Figure S2 (a), (b) & (c)). This score is reflective of the number of TRs binding within the CRM region. The total number of CRMs accessed from ReMap2022 and the ones retained post-filtering are summarized in Table 3.

**Table 3:**
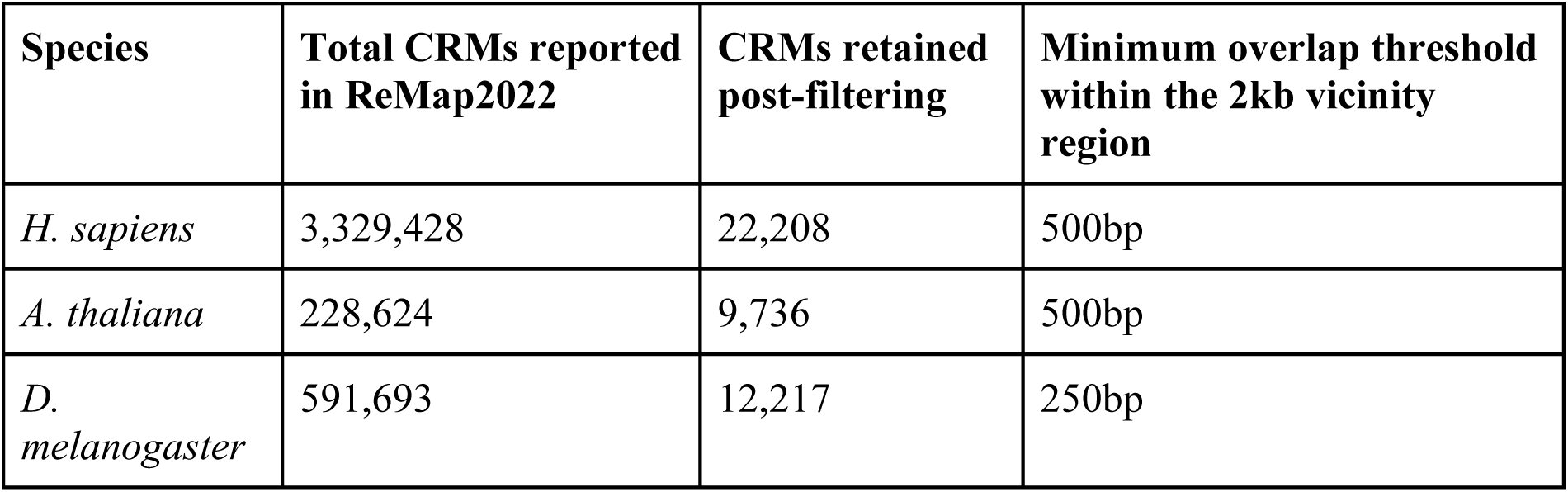
Summary of the CRM information retrieved from ReMap2022. The entire sets of CRMs were filtered to keep a subset which had a score of more than 30 and were in the 2kb vicinity of the coding regions. The threshold for the overlap with the vicinity is noted in the last column. The shorter overlap length for *D. melanogaster* (250bp) as compared to *H. sapiens* and *A. thaliana* (500bp) is explained by the comparatively compact nature of the *D. melanogaster* genome.

In addition to their genomic position, ReMap2022 also provides information on the TRs binding within each CRM. To identify the binding coordinates of each TR, we first retrieved the position weight matrix (PWM) information per TR from external databases (Table 4). Next, using *TEMPLE* (Litovchenko & Laurent, 2016), we scanned for potential TF-BS regions per CRM using the PWM information with a *p-value* cut-off of 0.05. Using *TEMPLE* we could identify the potential binding regions of each TRs within the CRMs given their binding profiles (Figure 1(a)).

**Figure 1.**
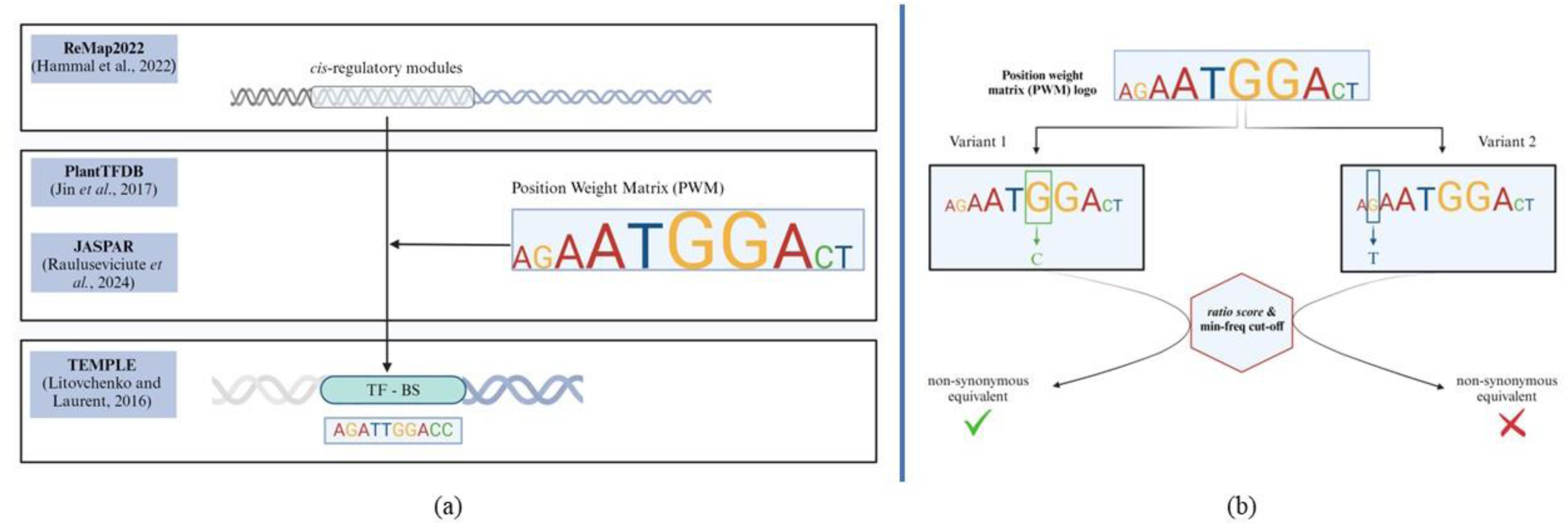
Identifying TF-BS regions and constructing the non-synonymous equivalent class within those regions. (a) – Using ReMap2022 we first extract the species-specific CRM regions, these regions are scanned for specific TF-BS regions by feeding the CRM coordinates and the PWM information of TRs to TEMPLE. (b) – The nonsynonymous equivalent class is identified using the ratio score (**see equation 1**) and frequency cutoff threshold. Variants having a ratio score above 0.6 and either of ancestral or derived frequency above 400 are shortlisted, the rest are not used in the analyses (Supplementary figure S3)

**Table 4:**
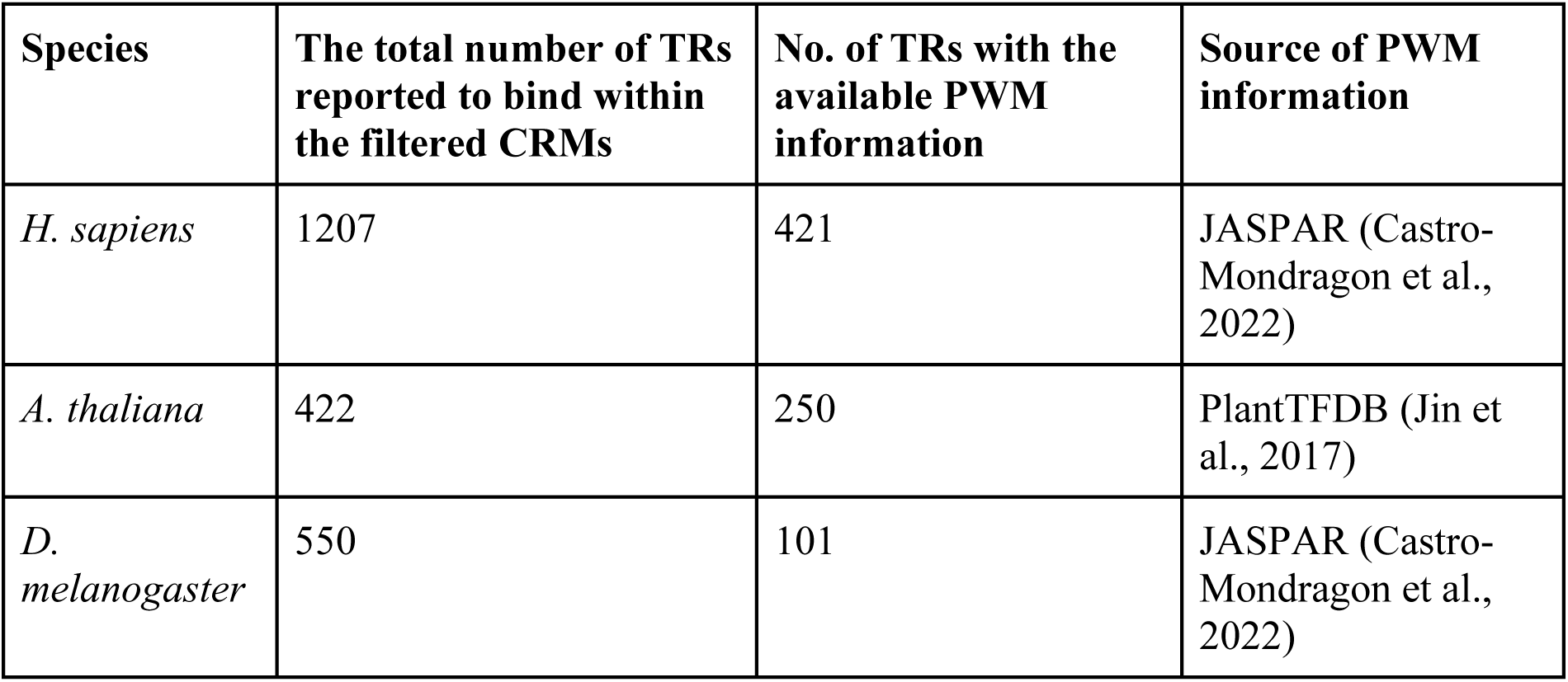
Summary of the identified TRs within the CRM regions and a subset of them used in this study which had accessible PWM information.

| Species | The total number of TRs reported to bind within the filtered CRMs | No. of TRs with the available PWM information | Source of PWM information |
| --- | --- | --- | --- |
| *H. sapiens* | 1207 | 421 | JASPAR (Castro-Mondragon et al., 2022) |
| *A. thaliana* | 422 | 250 | PlantTFDB (Jin et al., 2017) |
| *D. melanogaster* | 550 | 101 | JASPAR (Castro-Mondragon et al., 2022) |

### Constructing a non-synonymous equivalent class for TF-BS

One of the main challenges in performing a coding-region equivalent analysis of non-coding regions is to identify variants that are likely to have a functional consequence. In the case of coding regions, non-synonymous variants are perceived to have a functional consequence due to their impact on the encoded amino acid. To identify a class of non-synonymous equivalent variants in the non-coding TF-BS regions we used the PWM information. PWM contains position-specific count information per nucleotide representing the probability of the occurrence of each nucleotide per position. In addition to identifying the TF-BS regions, we also identified variants occurring within them. Using these counts and variant information, we investigated the change in PWM score between the variant and the ancestral allele. We captured this change in the PWM score of the ancestral and derived allele using a metric that we named the *ratio score*:

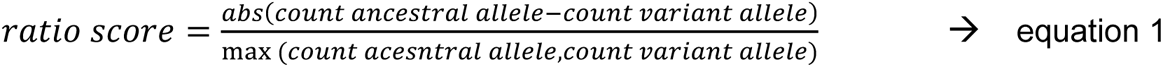

This metric captures the change in the counts for the ancestral and derived alleles. Variants that could impact the affinity would have a high *ratio score*. To minimize the inclusion of variants that would likely not impact the affinity, we set a *ratio score* cut-off of 0.6. Next, to eliminate the possibility of low counts influencing the identification of affinity-changing variants, we further filtered variants to keep only those whose ancestral or derived alleles had counts of 400 or more (Supplementary Figure S3). The variants retained post-filtering are shortlisted as non-synonymous equivalents for the TF-BS regions as they could potentially alter the binding affinity (Figure 1(b)).

Ascertaining an equivalent of the synonymous class through a similar approach is challenging due to the rigid cutoffs for segregating the variants. Specifically, variants not annotated as “non-synonymous” but having a ratio score and minimum counts close to the cutoffs may also impact the binding affinity. To overcome this, we used the synonymous variants from the coding regions (All genes) as a neutral class for the non-coding TF-BS regions as in previous studies (Kosakovsky Pond et al., 2005). Similar to the coding regions, the obtained proportions of non-synonymous variants (π_n_ and K_n_) were normalized by their respective Nei-Gojobori distances (Nei & Gojobori, 1986). Specifically, the total number of non-synonymous variants were normalized by the possible number of non-synonymous sites within the TF-BS regions. The total number of possible non-synonymous sites within the TF-BS were identified by similar cutoffs and filtering methods mentioned above.

### Calculation of α for the MK test of positive selection

The traditional MK test does not account for the presence of slightly deleterious variants segregating within species when calculating the proportion of adaptive substitutions, α. To account for this, tools like *asymptoticMK* (Haller & Messer, 2017) look at each mutational class in the Site-Frequency Spectrum (SFS) to make an improved estimate of α. Since this method estimates the α statistic per SFS class, it is sensitive to the number of variants present per class. Specifically, mainly due to short genomic lengths, the presence of a fewer number of variants in each class could result in inconsistent α estimates. Hence, the α estimates of comparatively shorter genomic regions (TF-BD, non-BD and TF-BS) could have high CIs. To account for this, we pooled the variants and employed a combination of the *asymptoticMK* and the traditional MK test. Specifically, we used the SFS class-specific distribution of the α estimates for the All genes class per species from *asymptoticMK* to decide on the low and high-frequency cutoffs and keep only the converged α values. Such cutoffs were placed by visual inspection on the convergence of the asymptote. This constituted a new dataset, from which we recalculated a single α value per genomic region using the traditional MK test. The CIs were constructed by performing 500 resampling cycles on the pooled polymorphism data.

### Simulation setup for different selfing regimes

To test the effect of selfing in the convergence of α we performed forward simulations using *SFScode* (Hernandez & Bateman, 2008). More specifically, we simulated four selfing regimes with selfing rates of 0.0, 0.3, 0.6, and 0.9. Each rate represents the proportion of individuals (in each generation) produced by selfing. Simulations included stretches of 1Mbp of coding DNA, with a mutation rate of 5e-09 per base pair, a recombination rate of 1e-08 per base pair, and a selective constraint of 2N_e_s=9, with N_e_=200000 (re-scaled to 500 for the forward simulations). We ran the simulations for 100000 generations, 45 individuals per population. Each simulation yielded FASTA files from which we used custom R scripts to count the number of synonymous and non-synonymous changes and calculated α for each SFS class and each selfing regime.

### Code availability

We developed two tools for performing the analyses of the genetic variants occurring within the coding and non-coding regions through a comparative and population genomics framework. *Alag* (https://github.com/Manaswwm/Alag) is tailored for performing the analyses of genetic variants occurring within the coding regions. In this study, we used *Alag* for the analyses of the TF-BD, non-BD and All genes regions. *templeRun* (https://github.com/Manaswwm/templeRun), a wrapper around TEMPLE (Litovchenko & Laurent, 2016), was developed to perform the analyses of genetic variants occurring within the TF-BS regions.

## Results

### Dataset and annotation of TFs

We compared genomic variation across TF-BDs and TF-BSs in three species: *Homo sapiens*, *Arabidopsis thaliana*, and *Drosophila melanogaster*. We used two populations per species, including populations from their ancestral geographical range. Samples were chosen in a way that minimized the effect of recent admixture or gene flow between the populations. For these three species we gathered genome-wide population genomic data, and a robust annotation of TFs (Swissprot) and cis-regulatory modules (Hammal et al., 2022). TFs were annotated based on SwissProt/InterPro and GO annotations. The species-specific number of TFs and the BDs identified within them are tabulated in Table 5. Using *Alag* we integrated the genome-wide annotations and variation occurring within the coding regions (TF-BD, non-BD and All regions).

**Table 5:** Summary the species-specific identified number of TFs and the corresponding number of TF-BDs.

| Species | No. of TFs | No. of BDs |
| --- | --- | --- |
| *H. sapiens* | 886 | 1198 |
| *A. thaliana* | 861 | 1030 |
| *D. melanogaster* | 217 | 325 |

### TF-BDs under high selective constraint at the population level

Comparing π_n_/π_s_ constraint ratios for all six populations and gene classes allowed us to make several comparative observations (Figure 2). First, the TF-BD regions seem to be under an increased constraint as compared to the two control regions. This signal was consistent across all populations. This suggests that, as compared to the two control regions, TF-BDs are more conserved which could be explained by their functional importance. Additionally, the relatively weak constraint exerted on non-BD in *A. thaliana* compared to *H. sapiens* and *D. melanogaster* represents an interesting observation. Next, in the context of the selection-drift balance, the high constraint values (depicted by low π_n_/π_s_) revealed that *D. melanogaster* experiences more negative selection than *H. sapiens* and *A. thaliana*. This higher amount of negative selection could be directly related to the larger N_e_ of *D. melanogaster* (∼1x10^6^, versus 2x10^5^ for *A. thaliana* and 2x10^4^ for humans). These observations are in agreement with the expectations under the drift-selection equilibria (Eyre-Walker & Keightley, 2007; Galtier, 2016; James et al., 2016).

**Figure 2.**
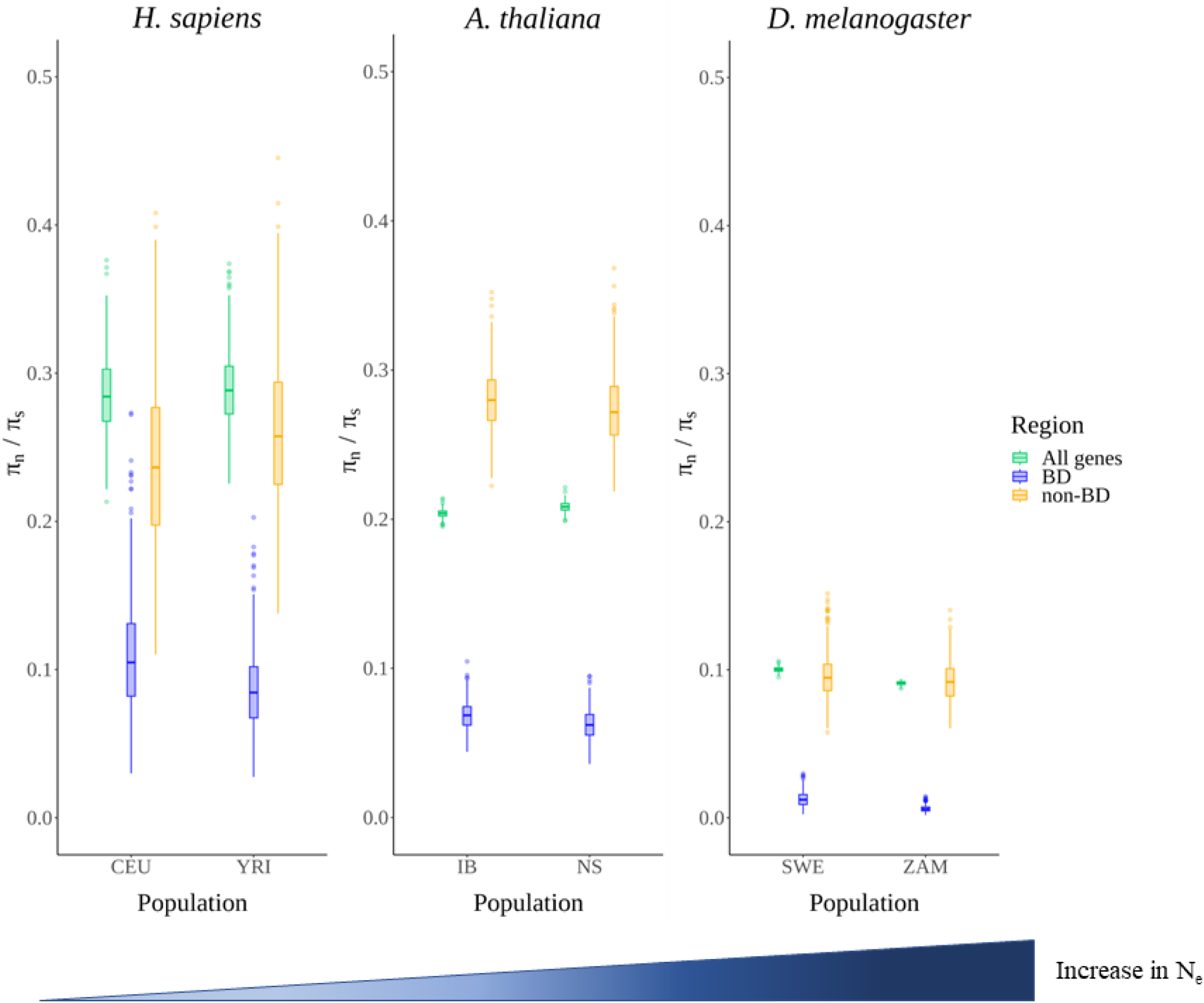
Comparisons of the coding region-specific π_n_/π_s_ constraint ratios across the six populations of the three species. Different colours represent the three classes of genomic regions. The distributions are constructed by performing 500 resampling cycles on the constraint ratios. The gradient blue triangle at the bottom is indicative of the overall increase in the N_e_. The population codes are: CEU – Utah residents with central European ancestry, YRI – Yoruba from Ibadan, IB – Iberia, NS – North Sweden, SWE – Sweden, ZAM – Zambia

### TF-BDs under high selective constraint across species

To find out whether these results obtained at the population level could be reproduced at the species level, we calculated constraint ratios (K_n_/K_s_) using pairwise genome-wide alignments between the three reference genomes of *H. sapiens*, *A. thaliana*, and *D. melanogaster* and their respective sister species *Pan troglodytes*, *Arabidopsis lyrata*, and *Drosophila simulans* (Figure 3). Our results showed that, as expected, the levels of constraint acting on TF-BDs are more extreme as compared to non-BD and All genes, thus the signal of high constraint acting on the TF-BDs was found to be consistent both at the population and species level. Interestingly, despite their vastly different N_e_, *H. sapiens* and *A. thaliana* had similar levels of K_n_/K_s_ as compared to *D. melanogaster*, who seem to have the lowest K_n_/K_s_ estimates. Hence, the relation between constraint and genetic drift was not detectable at the species level (Figure 3). While slightly deleterious variants tend to segregate more often and at higher frequencies in populations subject to significant levels of genetic drift, their chances of getting fixed in the whole species remain very low and comparable between species, irrespective of how much genetic drift they experience.

**Figure 3.**
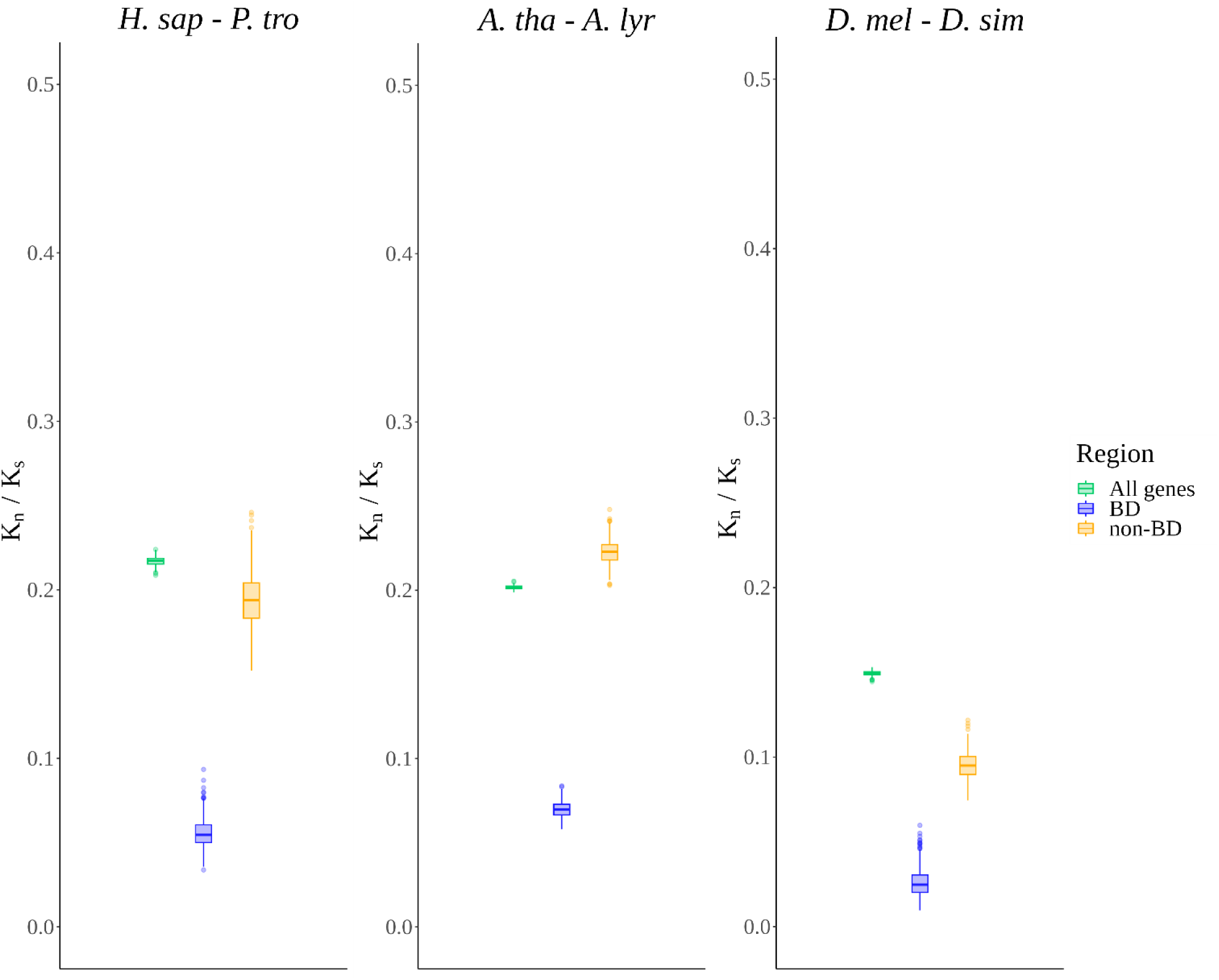
Comparison of the coding region-specific K_n_/K_s_ ratios between the three species and their respective outgroup species. Different colours represent the three classes of genomic regions. The distributions are constructed by performing 500 resampling cycles on the constraint ratios.

### TF-BDs harbour higher proportions of deleterious variants

Our analyses predict strong deleterious effects of mutations occurring in TF-BDs and, at the same time, weak efficiency of selection against these mutations in populations with high genetic drift (e.g. *H. sapiens*). Hence, we hypothesized that the naturally occurring variants within the TF-BD regions should be enriched for detrimental (pathogenic) variants. For this, we used the *ClinVar* dataset (Landrum et al., 2018), which catalogues variants within the human genome that have been highlighted to have clinical significance. Specifically, we compared the proportion of the pathogenic variants from the ClinVar database in the TF-BD and non-BD classes. We found that variants occurring within TF-BDs are significantly more likely to be detrimental to fitness than those occurring in non-BD regions (Fischer’s exact test, *p-value* < 0.01, Figure 4). This finding further supplements the comparatively high constraint observed on TF-BDs. Specifically, variants within TF-BDs could potentially disrupt multiple gene regulatory interactions, thereby having a detrimental impact on fitness.

**Figure 4.**
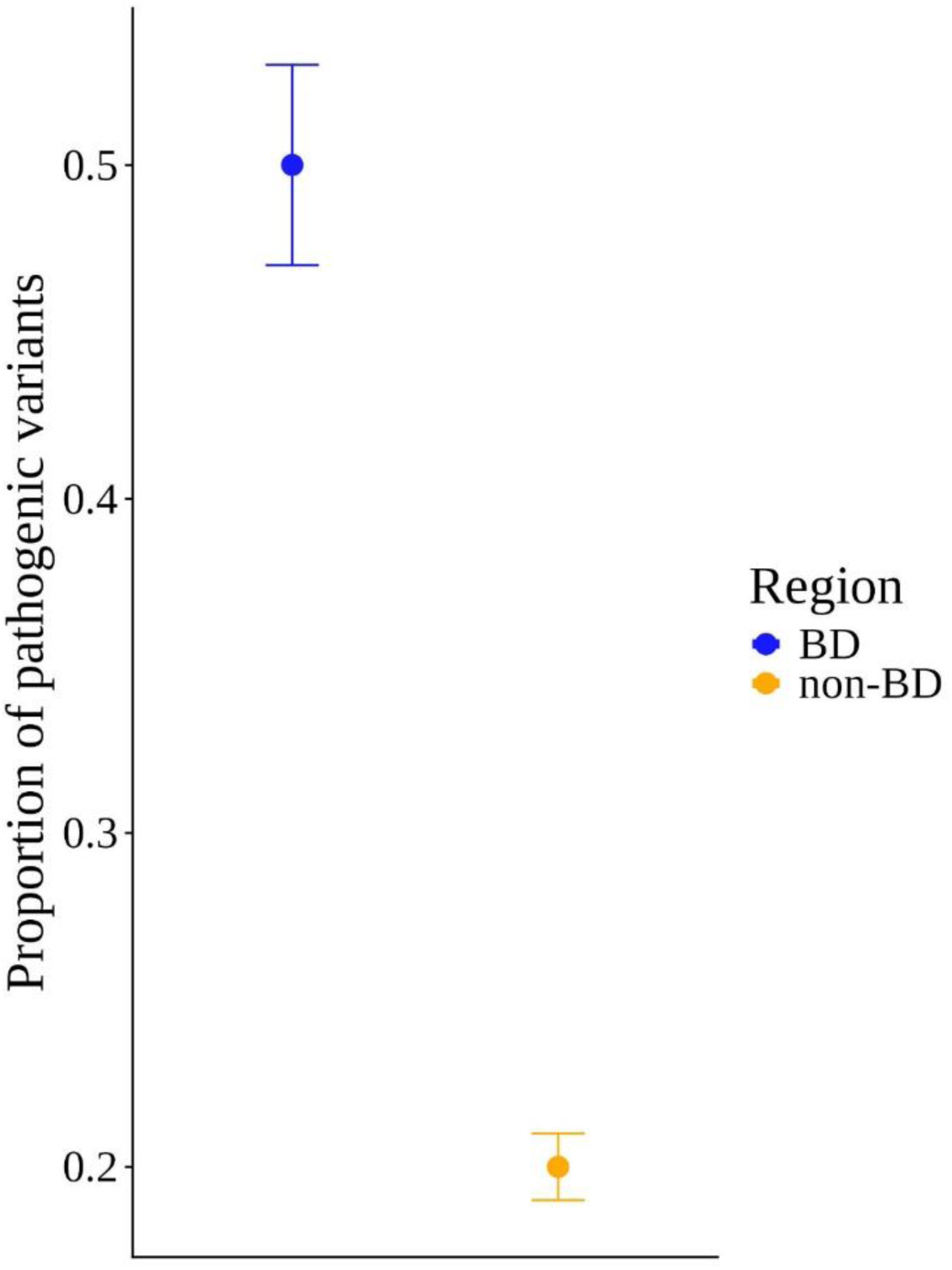
Comparing the proportions of pathogenic variants in the BD and non-BD regions – different colours indicate different regions (BD – blue, non-BD – yellow). The proportion of pathogenic variants is calculated by taking the per-region ratio of the total number of annotated pathogenic variants to the total sum of pathogenic and benign variants. The plot indicates the 95% CI with the bootstrapping method and the solid-coloured points indicate the mean.

### TF-BS regions under relaxed constraint when compared to coding regions

When comparing the distribution of the π_n_/π_s_ and the K_n_/K_s_ constraint ratios for the coding (All genes, TF-BD and non-BD) and non-coding TF-BS regions, we found that the non-coding regions are under an overall relaxed constraint when compared to the coding regions across both evolutionary timescales (Figure 5(a) and (b)). Interestingly, for the ancestral populations of large N_e_ species (IB for *A. thaliana* and ZAM for *D. melanogaster*), the constraint levels acting on the TF-BS regions were equivalent to their respective coding regions (*non-BD* for IB, *non-BD* and *All genes* for ZAM). Hence, even though TF-BSs are under a relaxed constraint as compared to the coding regions for small N_e_ species (*H. sapiens*), the constraint was observed to increase gradually with an increase in N_e_ and eventually reaches the levels of coding regions. This could be explained by the reduced drift for large N_e_ species, wherein selection acts on the non-coding functional elements on the levels comparable to the overall coding region. However, despite the reduction in drift, TF-BDs are still under higher constraints as compared to TF-BSs.

**Figure 5.**
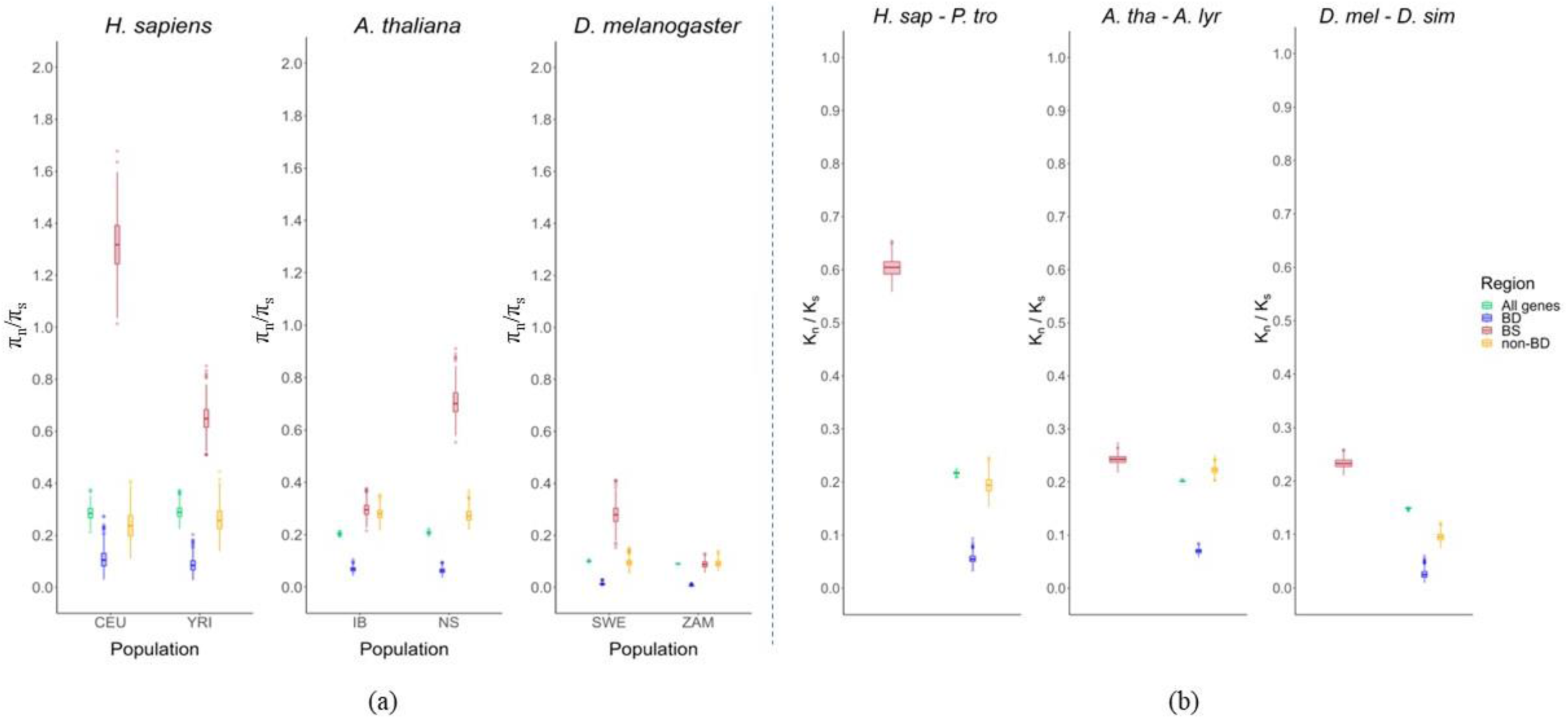
Comparing the constraint ratios for coding and non-coding regions. (a) – Comparing the π_n_/π_s_ constraint ratio & (b) – Comparing the K_n_/K_s_ constraint for TF-BS, TF-BD and the two control regions. The distributions are deduced by performing 500 resampling cycles. Different colours indicate the different genomic regions. The population codes are: CEU – Utah residents with central European ancestry, YRI – Yoruba from Ibadan, IB – Iberia, NS – North Sweden, SWE – Sweden, ZAM – Zambia

### Comparing species-specific estimates of adaptive evolution at transcription factors

To measure the intensity of positive selection, we calculated the α statistic using the *asymptoticMK* tool (Haller & Messer, 2017). In contrast to the traditional MK test, *asymptoticMK* makes use of the SFS information to account for the influence of slightly deleterious mutations in estimating the intensity of positive selection. First, we focused on elucidating the signals of positive selection in TF-BD regions and contrasting them to All genes and non-BD. For the All genes class, the intensity of positive selection increases with an increase in N_e_ (Supplementary Figure S4), which is visible by an increase in the α statistic. This result concurs also with the drift-selection equilibria observations from the negative selection analysis in the previous sections.

However, in the case of TF-BD and non-BD regions, we found a high variance in the α estimates, thus preventing us from drawing statistically significant observations. The reason for such high variance was the presence of comparatively fewer SNPs in SFS classes for those regions as compared to All genes. While the total number of variants collected in the three species for BD and non-BD regions is not small (Supplementary Table ST4), the *asymptoticMK* method requires, as an intermediate step, that α values be calculated for each variant-frequency class independently. Given that the number of variants decreases rapidly with increasing frequency classes, a large sampling variance is associated with most frequency classes in the calculation of α, resulting in large confidence intervals.

### Using customized frequency cut-offs per species for performing the traditional MK test

To address the issue of high variance around the α estimates for TF-BD and non-BD regions, we used a simpler calculation of α, which calculates it directly on the set of variants that have a frequency within frequency cut-off values (Charlesworth & Eyre-Walker, 2008). This cutoff is meant to exclude low-frequency classes, which could potentially be enriched for slightly deleterious non-synonymous variants. To define appropriate cut-off values for each species, we investigated the distribution of frequency-class-specific α values for All genes region produced by *asymptoticMK* (Figure 6) and manually defined cut-off values retaining frequency classes for which α values stabilized around their asymptote. Low-frequency cutoffs varied for each species.

**Figure 6.**
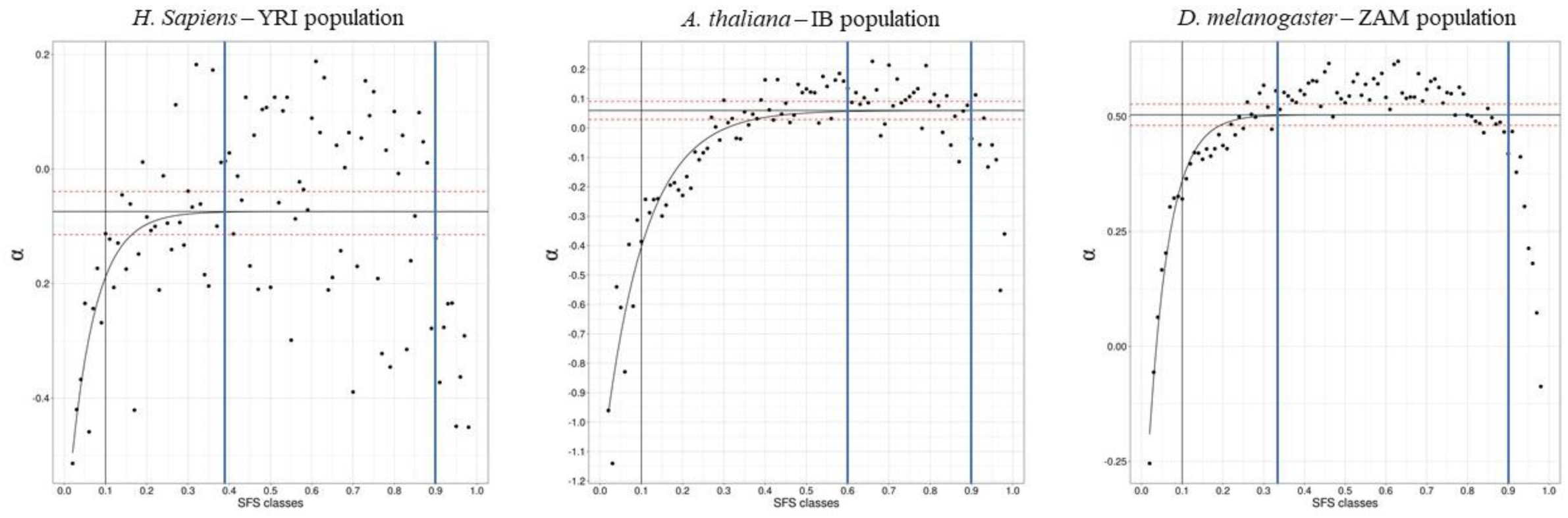
Species-specific asymptote convergence plots to estimate the α statistic using the *asymptoticMK* approach. The comparatively older populations per species were chosen as the representative population (*H. sapiens* – Yoruba from Ibadan, *A. thaliana* – Iberia and *D. melanogaster* – Zambia). The SFS classes are indicated on the X-axis and the SFS class-specific α estimates are indicated as solid black circles. The curved black lines indicate the asymptote and the convergence point indicates the α estimates (CI indicated with dotted horizontal red lines). The high- and low-frequency cutoffs are set by using the convergence of the asymptote as a reference. These cutoffs are indicated per species using solid blue vertical lines. The vertical black line is indicative of frequency 0.1, which is often used to remove the influence of slightly deleterious alleles.

We noted that after reaching a plateau at intermediate frequency classes, α values dropped again at high-frequency classes, which led us to exclude these frequency classes too. This is likely due to the well-documented presence of mis-polarized low-frequency variants among the highest frequency classes of the SFS when polarization is done using a single outgroup, as was the case in this study (Figure 6). High-frequency cutoffs were placed at a frequency of 0.9 for all three species. Following this procedure, we obtained corrected population-specific α values. After applying these steps and generating the new dataset, we observed that the highest α values come from *D. melanogaster* and intermediate values in *A. thaliana* (Figure 7). Particularly, we noted an overall increase in the α estimates for the regulatory TF-BD regions as compared to the other two control regions for species with larger N_e_ (*A. thaliana* and *D. melanogaster*). However, this signal fades for species with low N_e_ (*H. sapiens*). Overall, drift reduces the fixation probability of beneficial alleles, meaning that selection works best in large populations.

**Figure 7.**
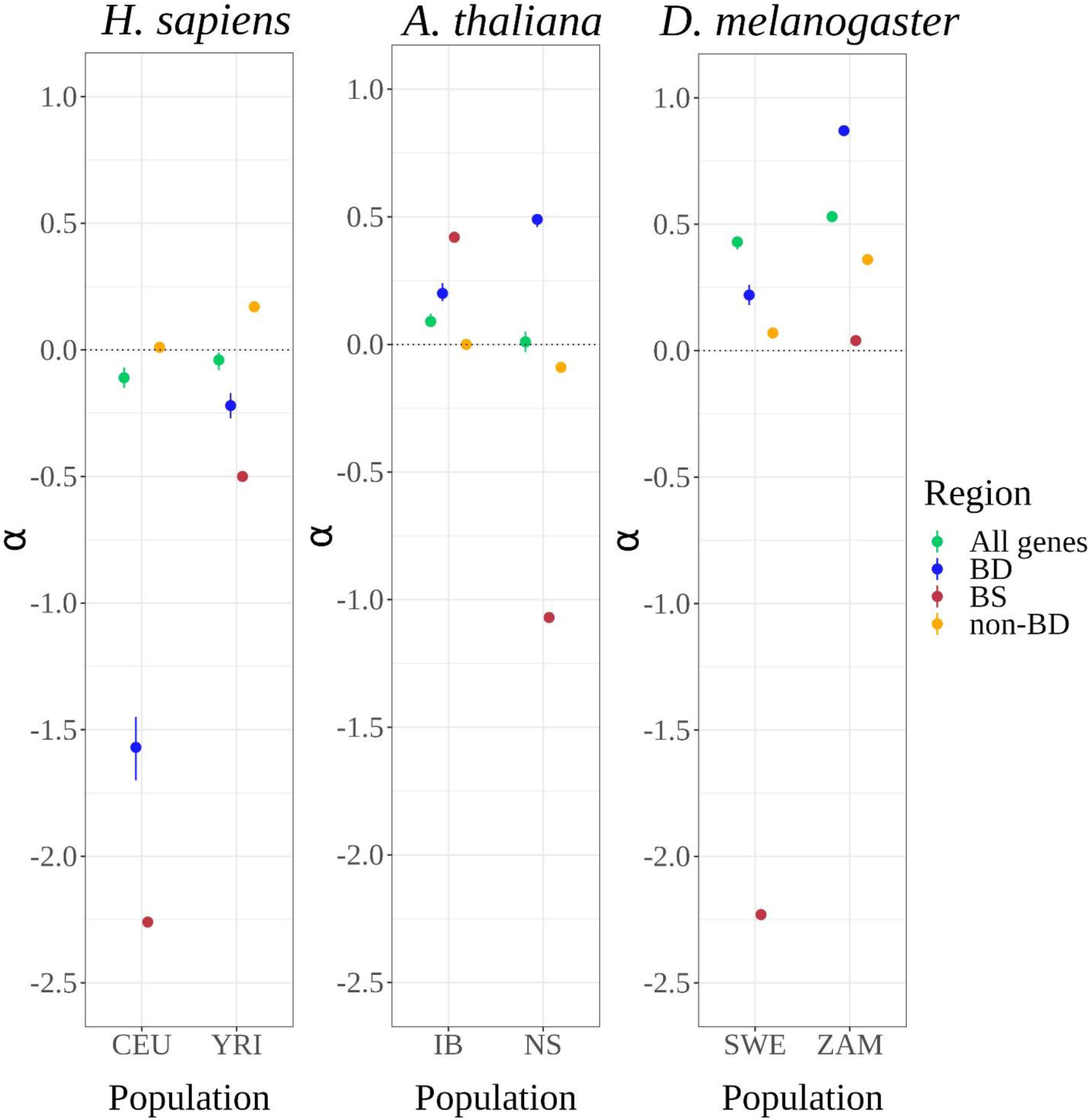
Estimating the coding and non-coding region-specific α using customized frequency cutoffs for the BD, BS and non-BD regions. On applying the species-specific frequency cutoffs (obtained from Figure 7), we pooled the variants and performed the traditional MK-test. The variance around the estimates is constructed by performing 500 resampling cycles of the polymorphism data. The α estimate for the All genes region is calculated using the *asymptoticMK* tool. Different colours indicate the four coding and noncoding regions. The population codes are: CEU – Utah residents with central European ancestry, YRI – Yoruba from Ibadan, IB – Iberia, NS – North Sweden, SWE – Sweden, ZAM – Zambia

## Discussion

This study focused on elucidating the action of natural selection acting on regulatory TF-DNA interactions. Given that these interactions are the initiation points for the transcription of the effector gene, this study focused on the two interacting motifs that play a vital role in these TF-DNA interactions: TF binding domains (TF-BDs) and TF binding sites (TF-BS). This is the first study investigating the regulatory evolution of both TF-BD and TF-BS elements through a population and comparative genomics framework (but see Joshi et al., 2021). This approach enabled us to observe whether the signals captured on the shorter evolutionary timescales (polymorphism) were robust and consistent on the longer evolutionary timescales (divergence) (Lawrie & Petrov, 2014). This study spanned three species, namely – *Homo sapiens*, *Arabidopsis thaliana* and *Drosophila melanogaster*, and six populations belonging to these species. The interaction domains were carefully identified using the available annotation on TF-BD (Bateman et al., 2023; Sigrist et al., 2002) and TF-BS regions (Castro-Mondragon et al., 2022; Jin et al., 2017). One of the limitations of this study is the conservative filters used to identify these interaction domains, which potentially reduced its number. Notably, the TFs belonging to many of the Zinc-Finger families were omitted due to our chosen annotation schema filters and the segmented nature of their binding domains (**Supplementary Table ST1, ST2 and ST3**). Nonetheless, the species-specific TFs are consistently selected using similar filters for all three species.

### Selection acting on TF-BDs

We found a signal of high constraint acting on the TF-BD regions when compared to the two control classes (non-BD and All genes). This signal was observed to be consistent across all six populations of the three species and both the evolutionary scales. The high constraint acting on binding domains could be attributed to the functional importance in the context of a gene regulatory network (GRN). Given the pleiotropic nature of TFs (Chesmore et al., 2016), TF-BDs interact with and control the expression of multiple downstream effector genes. Hence, variants within *TF-BDs* could disrupt multiple links within GRNs. This signal was further corroborated using clinical variants data from the ClinVar database (Landrum et al., 2018). In terms of annotated clinical variants, we found that TF-BDs harboured higher proportions of pathogenic variants when compared to the functionally non-annotated non-BD regions within the TFs, thus confirming the functional importance of the constraint.

### Constructing a non-synonymous equivalent for TF-BSs

Next, we aimed to analyze the signal of constraint acting on TF-BSs. To make the analyses on the TF-BS regions comparable to those of the coding regions, we constructed a non-synonymous equivalent class for TF-BSs. Specifically, we formulated a *ratio score* metric that scores every variant occurring within the TF-BS region to capture its impact on the binding affinity using the ancestral allele for polarization. Using conservative cutoffs, we identified a class of variants that is potentially the non-synonymous equivalent for the TF-BS regions. One of the limitations of this approach is that it is based on rigid cutoffs which may not accurately capture the binding dynamics of TF-DNA interactions. However, we employed conservative cutoffs to ensure the inclusion of variants that are more likely to impact binding affinities at the risk of losing variants that might also have an impact on the binding affinities (**see Materials and Methods**). The approach of annotating variants in the TF-BS regions based on their impact on the binding affinities has been performed before (He et al., 2011). However, this is the first study to apply it on the scale of multiple species and their respective populations.

### Selection acting on TF-BSs

The TF-BS regions from small N_e_ species (such as *H. sapiens*) were observed to be under relaxed constraint. The observation of non-coding elements being under relaxed constraint as compared to the coding elements is in line with several previous studies (Andolfatto, 2005; Haddrill et al., 2008; Naidoo et al., 2018; Torgerson et al., 2009). However, interestingly, we found that the levels of constraint on the TF-BS regions were comparable to that of the coding regions for the older populations of large N_e_ species (such as ZAM – *D. melanogaster* and IB – *A. thaliana*). In the case of the ZAM population, the level of constraint on the TF-BS region is similar to the overall protein-coding regions of the genome. These observations suggest that in species with reduced drift, natural selection acts with more efficiency also on non-coding regions. However, TF-BDs were consistently observed to be under higher constraints when compared to TF-BSs (Figure 5(a) and (b)).

One of the advantages of studying these three species was the differences in their respective effective population sizes (N_e_). It has been hypothesized that the efficiency of natural selection increases with an increase in the N_e_ due to the reduced influence of genetic drift (Eyre-Walker & Keightley, 2007; Galtier, 2016; James et al., 2016). The observations indicated that with an increase in N_e_, the efficiency of natural selection in removing non-beneficial and potentially detrimental alleles increases, thereby resulting in reduced π_n_/π_s_ ratios. More precisely, on comparing the π_n_/π_s_ constraint ratios for the four genomic regions (TF-BD, TF-BS, non-BD and All genes) across the populations of all three species, we found a decrease in π_n_/π_s_ in parallel to an increase in N_e_. This observation was in line with the N_e_ hypothesis mentioned before. Particularly, we observed lower π_n_/π_s_ ratios for species with high N_e_ (*D. melanogaster*) when compared to species with low N_e_ (*H. sapiens*). In the case of TF-BS regions, this gradient was even seen for populations within the same species (Figure 5(a)). Specifically, we noted a higher π_n_/π_s_ ratio for the populations with comparatively lower drift (*YRI – H. sapiens*, *IB – A. thaliana* and *ZAM – D. melanogaster*) when compared to populations with comparatively higher drift levels (*CEU – H .sapiens*, *NS – A. thaliana* and *SWE* – *D. melanogaster*). We notice however that, in *A. thaliana*, π_n_/π_s_ values in the All genes class were lower than in the non-BD class.

### Quantifying positive selection through α

Next, to quantify the action of positive selection, we developed a method derived from the asymptoticMK approach (Haller & Messer, 2017) for regions with small number of SNP information. We proposed this method to accurately identify the frequency cutoff to limit the influence of slightly deleterious variants segregating within the species. On using *asymptoticMK* for TF-BD and non-BD regions, we noted a high variance around the α estimate which could be explained by the relatively lower number of SNPs per SFS classes. For such regions with fewer variants (TF-BD, TF-BS and non-BD) we used the convergence point of the asymptote to determine the frequency cutoff per species. On pooling the retained variants, we estimated the intensity of positive selection using the α statistic. Here, we found weak signals of high α for TF-BD regions when compared to the other classes for high N_e_ species (*A. thaliana* and *D. melanogaster*), suggesting positive selection might also act with increased efficiency on TF-BDs. However, these signals were not as decisive as the signals for high constraint on the TF-BD regions.

### Selfing

We found that the α estimates from *asymptoticMK* for the All genes region also showed a gradient of increase with an increase in N_e_. This observation is also in agreement with the N_e_ hypothesis (Galtier, 2016). Specifically, positive selection acts with higher efficiency on large N_e_ species due to the reduction in genetic drift. However, the convergence points of the asymptote were not in line with the increase in N_e_. Specifically, in the case of *A. thaliana*, the convergence of the α asymptote occurred on comparatively higher frequencies as compared to the other two species (Figure 6). This could be explained by the influence of slightly deleterious variants in higher frequencies. One of the factors contributing to the presence of slightly deleterious variants in higher frequencies could be the mating system of the species. Of the three species, *A. thaliana* is a predominantly selfing plant species (Bechsgaard et al., 2006), while the other two (*H. sapiens* and *D. melanogaster*) rely solely on sexual reproduction. Due to their selfing nature, the genomes of selfing species display lower recombination rates which consequently hampers the ability of natural selection to remove the potentially deleterious mutations from the genome. This limitation is absent in the sexually reproducing species due to the crossing-over of chromosomes during meiosis. To confirm if differential selfing regimes influence the convergence of the asymptote, we performed forward simulations, which proved this hypothesis (Supplementary Figure S5).

This study probed the impact of natural selection acting on the regulatory motifs involved in the regulatory TF-DNA interactions through a comparative framework. This comparative framework enabled us to make comparisons across multiple species using commonly used summary statistics. The high constraint on TF-BD regions is expected given their functional importance, and this was further corroborated using empirical clinical variant data. Next, using a binding-affinity based approach, we aimed to compare the impact of natural selection on the non-coding TF-BS regions. We report that TF-BS are under relaxed constraint for small N_e_ species, however, their levels of constraint become comparable to that of coding regions for large N_e_ species. This finding suggests that with a reduction in drift, selection acts with higher intensity on not only the coding but also the functional non-coding elements. Finally, the signals for positive selection seemed to overall follow a similar trend as that of negative selection albeit with less intensity. Overall, our findings across all genomic regions are also in concurrence with the overall drift-selection equilibria.

## Acknowledgements

We thank Miltos Tsiantis for supporting this work through a Max Planck Society core grant to the Department of Comparative Development and Genetics.

## Notes

### Competing Interest Statement

The authors have declared no competing interest.

## References

Altschul, S. F., Gish, W., Miller, W., Myers, E. W., & Lipman, D. J. (1990). Basic local alignment search tool. Journal of Molecular Biology, 215(3), 403–410. 10.1016/S0022-2836(05)80360-2

Andolfatto, P. (2005). Adaptive evolution of non-coding DNA in Drosophila. Nature, 437(7062), 1149–1152. 10.1038/nature04107

Arbiza, L., Gronau, I., Aksoy, B. A., Hubisz, M. J., Gulko, B., Keinan, A., & Siepel, A. (2013). Genome-wide inference of natural selection on human transcription factor binding sites. Nat Genet, 45(7), 723–729. 10.1038/ng.2658

Bateman, A., Martin, M. J., Orchard, S., Magrane, M., Ahmad, S., Alpi, E., Bowler-Barnett, E. H., Britto, R., Bye-A-Jee, H., Cukura, A., Denny, P., Dogan, T., Ebenezer, T. G., Fan, J., Garmiri, P., da Costa Gonzales, L. J., Hatton-Ellis, E., Hussein, A., Ignatchenko, A., … Zhang, J. (2023). UniProt: the Universal Protein Knowledgebase in 2023. Nucleic Acids Research, 51(D1), D523–D531. 10.1093/NAR/GKAC1052

Bechsgaard, J. S., Castric, V., Charlesworth, D., Vekemans, X., & Schierup, M. H. (2006). The Transition to Self-Compatibility in Arabidopsis thaliana and Evolution within S-Haplotypes over 10 Myr. Molecular Biology and Evolution, 23(9), 1741– 1750. 10.1093/MOLBEV/MSL042

Castro-Mondragon, J. A., Riudavets-Puig, R., Rauluseviciute, I., Berhanu Lemma, R., Turchi, L., Blanc-Mathieu, R., Lucas, J., Boddie, P., Khan, A., Perez, N. M., Fornes, O., Leung, T. Y., Aguirre, A., Hammal, F., Schmelter, D., Baranasic, D., Ballester, B., Sandelin, A., Lenhard, B., … Mathelier, A. (2022). JASPAR 2022: the 9th release of the open-access database of transcription factor binding profiles. Nucleic Acids Research, 50(D1), D165–D173. 10.1093/NAR/GKAB1113

Charlesworth, J., & Eyre-Walker, A. (2008). The McDonald-Kreitman test and slightly deleterious mutations. Molecular Biology and Evolution, 25(6), 1007–1015. 10.1093/molbev/msn005

Chesmore, K. N., Bartlett, J., Cheng, C., & Williams, S. M. (2016a). Complex Patterns of Association between Pleiotropy and Transcription Factor Evolution. 8(10), 3159–3170. 10.1093/GBE/EVW228

Chesmore, K. N., Bartlett, J., Cheng, C., & Williams, S. M. (2016b). Complex Patterns of Association between Pleiotropy and Transcription Factor Evolution. Genome Biology and Evolution, 8(10), 3159–3170. 10.1093/GBE/EVW228

Connelly, C. F., Skelly, D. A., Dunham, M. J., & Akey, J. M. (2013). Population genomics and transcriptional consequences of regulatory motif variation in globally diverse saccharomyces cerevisiae strains. Molecular Biology and Evolution, 30(7), 1605–1613. 10.1093/molbev/mst073

Desvergne, B., Michalik, L., & Wahli, W. (2006). Transcriptional regulation of metabolism. 86(2), 465–514. https://pubmed.ncbi.nlm.nih.gov/16601267/

Edgar, R. C. (2004). MUSCLE: multiple sequence alignment with high accuracy and high throughput. Nucleic Acids Research, 32(5), 1792. 10.1093/NAR/GKH340

Ellegren, H. (2008). Comparative genomics and the study of evolution by natural selection. Molecular Ecology, 17(21), 4586–4596. 10.1111/J.1365-294X.2008.03954.X

Eyre-Walker, A., & Keightley, P. D. (2007). The distribution of fitness effects of new mutations. Nature Reviews Genetics, 8(8), 610–618. 10.1038/nrg2146

Galtier, N. (2016). Adaptive Protein Evolution in Animals and the Effective Population Size Hypothesis. PLOS Genetics, 12(1), e1005774. 10.1371/JOURNAL.PGEN.1005774

Haddrill, P. R., Bachtrog, D., & Andolfatto, P. (2008). Positive and negative selection on noncoding DNA in Drosophila simulans. Molecular Biology and Evolution, 25(9), 1825–1834. 10.1093/molbev/msn125

Haller, B. C., & Messer, P. W. (2017). AsymptoticMK: A web-based tool for the asymptotic McDonald-Kreitman test. G3: *Genes, Genomes, Genetics*, *7*(5), 1569–1575. 10.1534/g3.117.039693

Hammal, F., De Langen, P., Bergon, A., Lopez, F., & Ballester, B. (2022). ReMap 2022: a database of Human, Mouse, Drosophila and Arabidopsis regulatory regions from an integrative analysis of DNA-binding sequencing experiments. Nucleic Acids Research, 50(D1), D316–D325. 10.1093/NAR/GKAB996

He, B. Z., Holloway, A. K., Maerkl, S. J., & Kreitman, M. (2011). Does positive selection drive transcription factor binding site turnover? a test with drosophila cis-regulatory modules. PLoS Genetics, 7(4). 10.1371/journal.pgen.1002053

Hernandez, R. D., & Bateman, A. (2008). A flexible forward simulator for populations subject to selection and demography. Bioinformatics, 24(23), 2786–2787. 10.1093/BIOINFORMATICS/BTN522

James, J., Castellano, D., & Eyre-Walker, A. (2016). DNA sequence diversity and the efficiency of natural selection in animal mitochondrial DNA. Heredity, 118, 88–95. 10.1038/hdy.2016.108

Jin, J., Tian, F., Yang, D. C., Meng, Y. Q., Kong, L., Luo, J., & Gao, G. (2017). PlantTFDB 4.0: toward a central hub for transcription factors and regulatory interactions in plants. Nucleic Acids Research, 45(D1), D1040–D1045. 10.1093/NAR/GKW982

Joshi, M., Kapopoulou, A., & Laurent, S. (2021). Impact of Genetic Variation in Gene Regulatory Sequences: A Population Genomics Perspective. Frontiers in Genetics, 12(July), 1–10. 10.3389/fgene.2021.660899

Kosakovsky Pond, S. L., Frost, S. D. W., & Muse, S. V. (2005). HyPhy: Hypothesis testing using phylogenies. Bioinformatics, 21(5), 676–679. 10.1093/bioinformatics/bti079

Landrum, M. J., Lee, J. M., Benson, M., Brown, G. R., Chao, C., Chitipiralla, S., Gu, B., Hart, J., Hoffman, D., Jang, W., Karapetyan, K., Katz, K., Liu, C., Maddipatla, Z., Malheiro, A., McDaniel, K., Ovetsky, M., Riley, G., Zhou, G., … Maglott, D. R. (2018). ClinVar: improving access to variant interpretations and supporting evidence. Nucleic Acids Research, 46(D1), D1062–D1067. 10.1093/NAR/GKX1153

Lawrie, D. S., & Petrov, D. A. (2014). Comparative population genomics: power and principles for the inference of functionality. Trends in Genetics, 30(4), 133–139. 10.1016/J.TIG.2014.02.002

Lee, T. I., & Young, R. A. (2013). Transcriptional regulation and its misregulation in disease. Cell, 152(6), 1237–1251. https://pubmed.ncbi.nlm.nih.gov/23498934/

Litovchenko, M., & Laurent, S. (2016). TEMPLE: analysing population genetic variation at transcription factor binding sites. Molecular Ecology Resources, 16(6), 1428–1434. 10.1111/1755-0998.12535

Morales, J., Pujar, S., Loveland, J. E., Astashyn, A., Bennett, R., Berry, A., Cox, E., Davidson, C., Ermolaeva, O., Farrell, C. M., Fatima, R., Gil, L., Goldfarb, T., Gonzalez, J. M., Haddad, D., Hardy, M., Hunt, T., Jackson, J., Joardar, V. S., … Murphy, T. D. (2022). A joint NCBI and EMBL-EBI transcript set for clinical genomics and research. Nature 2022 *604*:7905, *604*(7905), 310–315. 10.1038/s41586-022-04558-8

Mu, X. J., Lu, Z. J., Kong, Y., Lam, H. Y. K., & Gerstein, M. B. (2011). Analysis of genomic variation in non-coding elements using population-scale sequencing data from the 1000 Genomes Project. Nucleic Acids Research, 39(16), 7058– 7076. 10.1093/nar/gkr342

Naidoo, T., Sjödin, P., Schlebusch, C., & Jakobsson, M. (2018). Patterns of variation in cis-regulatory regions: Examining evidence of purifying selection. BMC Genomics, 19(1), 1–14. 10.1186/s12864-017-4422-y

Nei, M., & Gojobori, T. (1986). Simple methods for estimating the numbers of synonymous and nonsynonymous nucleotide substitutions. Molecular Biology and Evolution, 3(5), 418–426. 10.1093/OXFORDJOURNALS.MOLBEV.A040410

Parsch, J., Novozhilov, S., Saminadin-Peter, S. S., Wong, K. M., & Andolfatto, P. (2010). On the Utility of Short Intron Sequences as a Reference for the Detection of Positive and Negative Selection in Drosophila. Molecular Biology and Evolution, 27(6), 1226–1234. 10.1093/MOLBEV/MSQ046

Romani, F., & Moreno, J. E. (2021). Molecular mechanisms involved in functional macroevolution of plant transcription factors. New Phytologist, 230(4), 1345– 1353. 10.1111/NPH.17161

Sigrist, C. J. A., Cerutti, L., Hulo, N., Gattiker, A., Falquet, L., Pagni, M., Bairoch, A., & Bucher, P. (2002). PROSITE: A documented database using patterns and profiles as motif descriptors. Briefings in Bioinformatics, 3(3), 265–274. 10.1093/BIB/3.3.265

Torgerson, D. G., Boyko, A. R., Hernandez, R. D., Indap, A., Hu, X., White, T. J., Sninsky, J. J., Cargill, M., Adams, M. D., Bustamante, C. D., & Clark, A. G. (2009). Evolutionary processes acting on candidate cis-regulatory regions in humans inferred from patterns of polymorphism and divergence. PLoS Genetics, 5(8). 10.1371/journal.pgen.1000592

Vernot, B., Stergachis, A. B., Maurano, M. T., Vierstra, J., Neph, S., Thurman, R. E., Stamatoyannopoulos, J. A., & Akey, J. M. (2012). Personal and population genomics of human regulatory variation. Genome Research, 22(9), 1689–1697. 10.1101/gr.134890.111

Vitti, J. J., Grossman, S. R., & Sabeti, P. C. (2013). Detecting natural selection in genomic data. Annual Review of Genetics, 47, 97–120. 10.1146/ANNUREV-GENET-111212-133526

Alonso-Blanco, C., Andrade, J., Becker, C., Bemm, F., Bergelson, J., Borgwardt, K. M. M., Cao, J., Chae, E., Dezwaan, T. M. M., Ding, W., Ecker, J. R. R., Exposito-Alonso, M., Farlow, A., Fitz, J., Gan, X., Grimm, D. G. G., Hancock, A. M. M., Henz, S. R. R., Holm, S., … Zhou, X. (2016). 1,135 Genomes Reveal the Global Pattern of Polymorphism in Arabidopsis thaliana. Cell, 166(2), 481–491. 10.1016/j.cell.2016.05.063

Auton, A., Abecasis, G. R., Altshuler, D. M., Durbin, R. M., Bentley, D. R., Chakravarti, A., Clark, A. G., Donnelly, P., Eichler, E. E., Flicek, P., Gabriel, S. B., Gibbs, R. A., Green, E. D., Hurles, M. E., Knoppers, B. M., Korbel, J. O., Lander, E. S., Lee, C., Lehrach, H., … Schloss, J. A. (2015). A global reference for human genetic variation. Nature, 526(7571), 68–74. 10.1038/nature15393

Byrska-Bishop, M., Evani, U. S., Zhao, X., Basile, A. O., Abel, H. J., Regier, A. A., Corvelo, A., Clarke, W. E., Musunuri, R., Nagulapalli, K., Fairley, S., Runnels, A., Winterkorn, L., Lowy, E., Eichler, E. E., Korbel, J. O., Lee, C., Marschall, T., Devine, S. E., … Zody, M. C. (2022). High-coverage whole-genome sequencing of the expanded 1000 Genomes Project cohort including 602 trios. Cell, 185(18), 3426–3440.e19. 10.1016/J.CELL.2022.08.004

Clark, A. G., Eisen, M. B., Smith, D. R., Bergman, C. M., Oliver, B., Markow, T. A., Kaufman, T. C., Kellis, M., Gelbart, W., Iyer, V. N., Pollard, D. A., Sackton, T. B., Larracuente, A. M., Singh, N. D., Abad, J. P., Abt, D. N., Adryan, B., Aguade, M., Akashi, H., … MacCallum, I. (2007). Evolution of genes and genomes on the Drosophila phylogeny. Nature 2007 *450*:7167, *450*(7167), 203–218. 10.1038/nature06341

Cunningham, F., Allen, J. E., Allen, J., Alvarez-Jarreta, J., Amode, M. R., Armean, I. M., Austine-Orimoloye, O., Azov, A. G., Barnes, I., Bennett, R., Berry, A., Bhai, J., Bignell, A., Billis, K., Boddu, S., Brooks, L., Charkhchi, M., Cummins, C., Da Rin Fioretto, L., … Flicek, P. (2022). Ensembl 2022. Nucleic Acids Research, 50(D1), D988–D995. 10.1093/NAR/GKAB1049

Hu, T. T., Pattyn, P., Bakker, E. G., Cao, J., Cheng, J. F., Clark, R. M., Fahlgren, N., Fawcett, J. A., Grimwood, J., Gundlach, H., Haberer, G., Hollister, J. D., Ossowski, S., Ottilar, R. P., Salamov, A. A., Schneeberger, K., Spannagl, M., Wang, X., Yang, L., … Guo, Y. L. (2011). The Arabidopsis lyrata genome sequence and the basis of rapid genome size change. Nature Genetics, 43(5), 476–483. 10.1038/NG.807

Kapopoulou, A., Kapun, M., Pieper, B., Pavlidis, P., Wilches, R., Duchen, P., Stephan, W., & Laurent, S. (2020). Demographic analyses of a new sample of haploid genomes from a Swedish population of Drosophila melanogaster. 10(1), 1–8. https://pubmed.ncbi.nlm.nih.gov/33376238/

Lack, J. B., Cardeno, C. M., Crepeau, M. W., Taylor, W., Corbett-Detig, R. B., Stevens, K. A., Langley, C. H., & Pool, J. E. (2015). The Drosophila genome nexus: a population genomic resource of 623 Drosophila melanogaster genomes, including 197 from a single ancestral range population. Genetics, 199(4), 1229–1241. 10.1534/GENETICS.115.174664

Mikkelsen, T. S., Hillier, L. W., Eichler, E. E., Zody, M. C., Jaffe, D. B., Yang, S. P., Enard, W., Hellmann, I., Lindblad-Toh, K., Altheide, T. K., Archidiacono, N., Bork, P., Butler, J., Chang, J. L., Cheng, Z., Chinwalla, A. T., Dejong, P., Delehaunty, K. D., Fronick, C. C., … Waterston, R. H. (2005). Initial sequence of the chimpanzee genome and comparison with the human genome. Nature 2005 *437*:7055, *437*(7055), 69–87. 10.1038/nature04072

