## Supplementary figures for "Comparative and population genomics analyses of the regulatory Transcription Factor (TF)-DNA interaction"

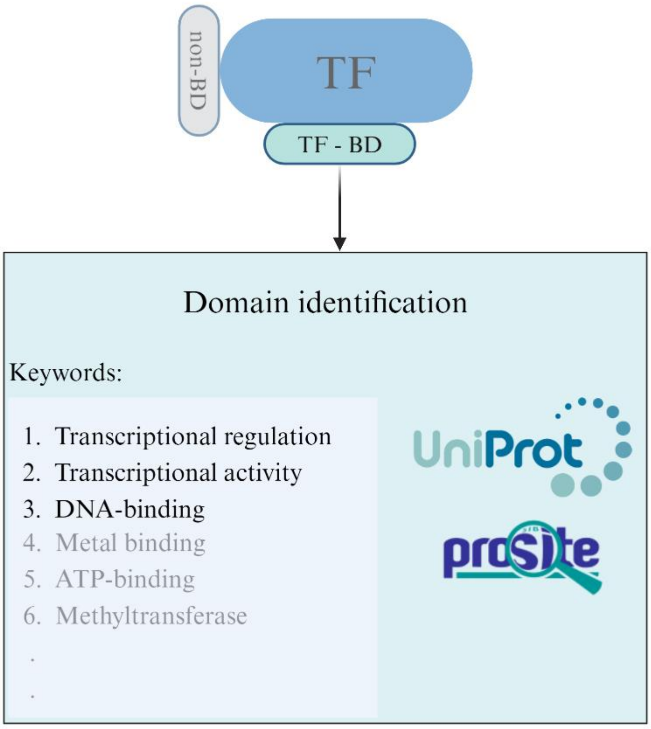


Figure S1: Identifying the TF-BD regions using the ontology terms. The annotated domains within the representative UniProt transcript are scanned for the following keywords – “Transcriptional regulation”, “Transcriptional activity” and “DNA-binding”. Domains whose annotations match these criteria are shortlisted as BD.

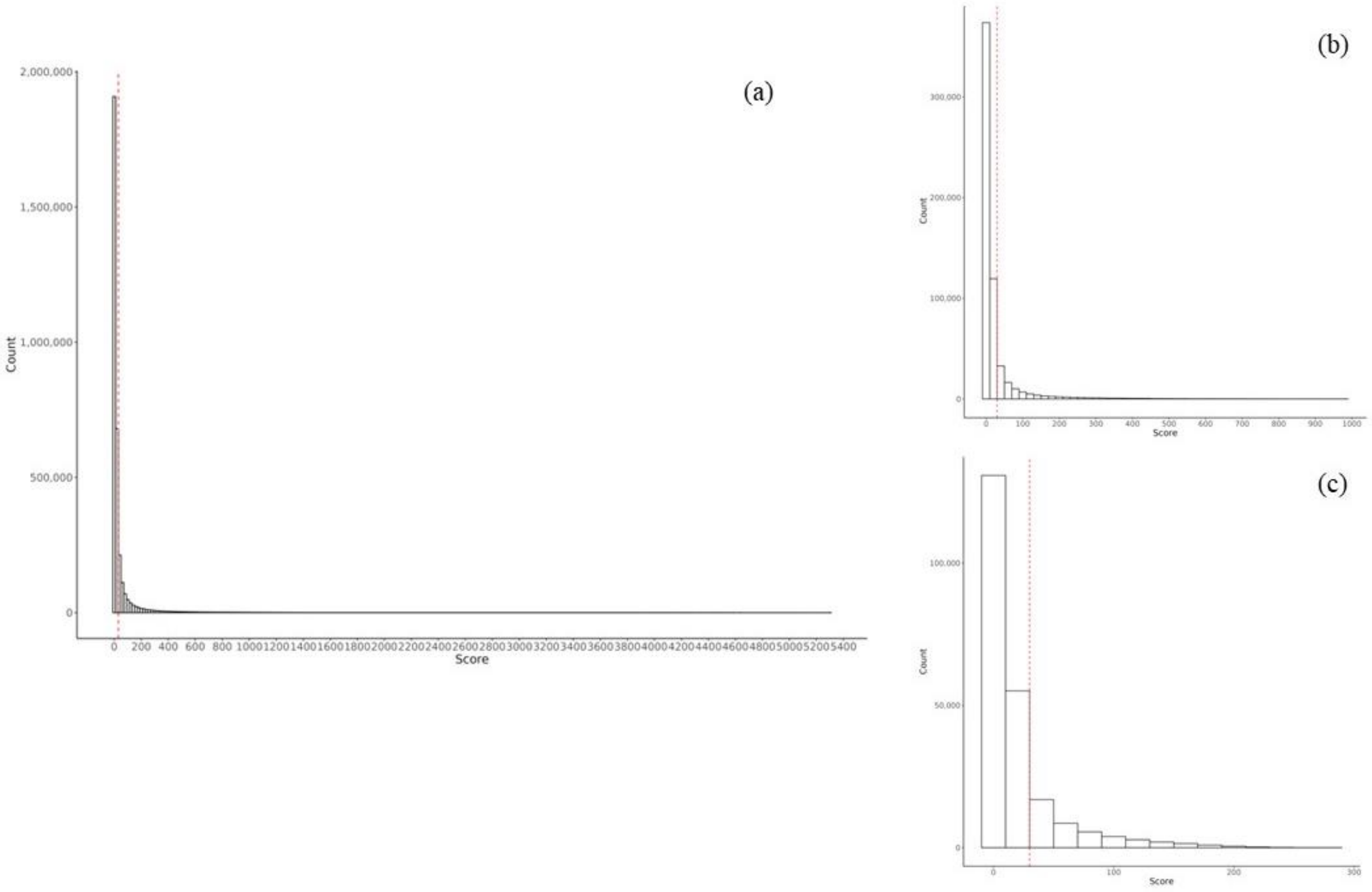


Figure S2: The species-specific CRM score distribution, here the score is indicated on the X-axis and the bar indicates the number of CRMs having the respective score. The scores are reflective of the number of TRs showing binding activity within the CRMs. The cut-off score is set at 30 to exclude regions to which a relatively smaller number of TRs are annotated to bind, this is done to obtain a more refined set of CRMs. The cutoff is shown with red dotted lines, the CRMs to the right of the cutoff are included in this study (score > 30), while those on the left are excluded. The three panels are indicative of the three species – (a) – *H. sapiens*; (b) – *A. thaliana*; (c) – *D. melanogaster*

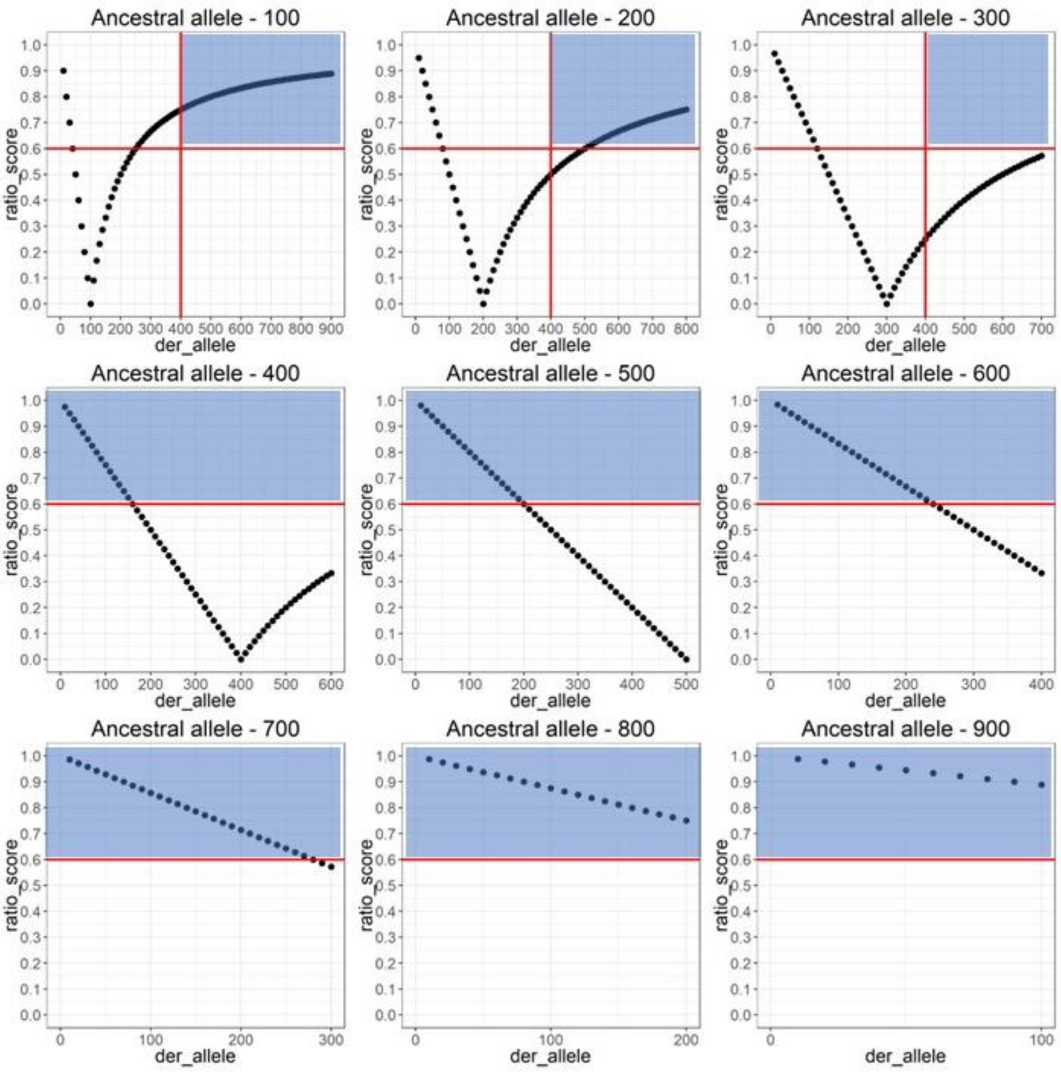


Figure S3: Subsetting the non-synonymous equivalent using the ratio score and frequency cutoff thresholds in different ancestral and derived allele count combinations. Here, every panel indicates a specific count of the ancestral allele (in step 100), the x-axis indicates the derived allele counts (in step 10) and the y-axis indicates the ratio score. The horizontal and vertical red lines indicate the ratio score and frequency cutoffs respectively. The blue boxes highlight the subset of variants that would be annotated as nonsynonymous equivalents.

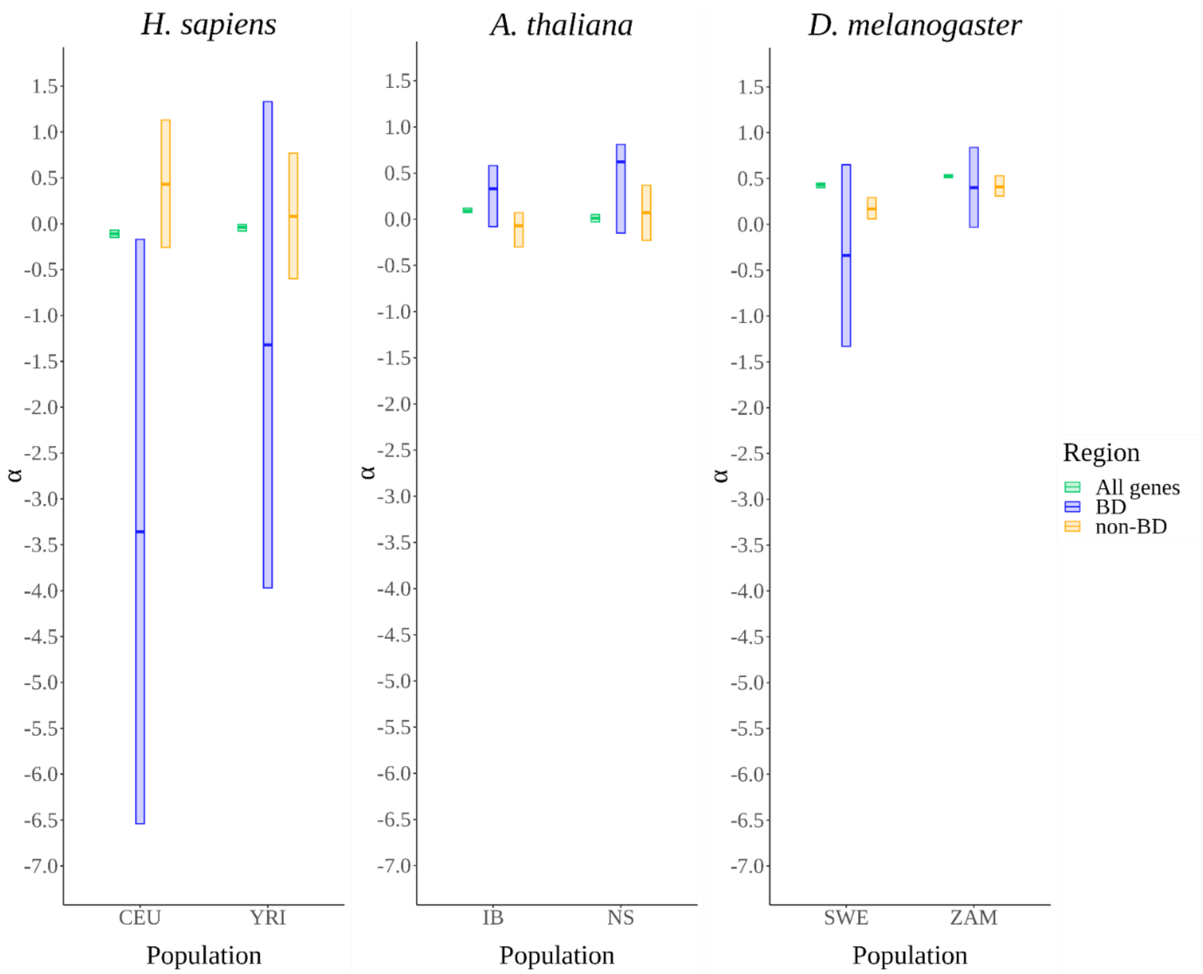


Figure S4 – Distribution of the α statistic estimates using the *asymptoticMK* tool for the coding regions across six populations. Different colours indicate the three different coding regions. The population codes are: CEU – Utah residents with central European ancestry, YRI – Yoruba from Ibadan, IB – Iberia, NS – North Sweden, SWE – Sweden, ZAM – Zambia

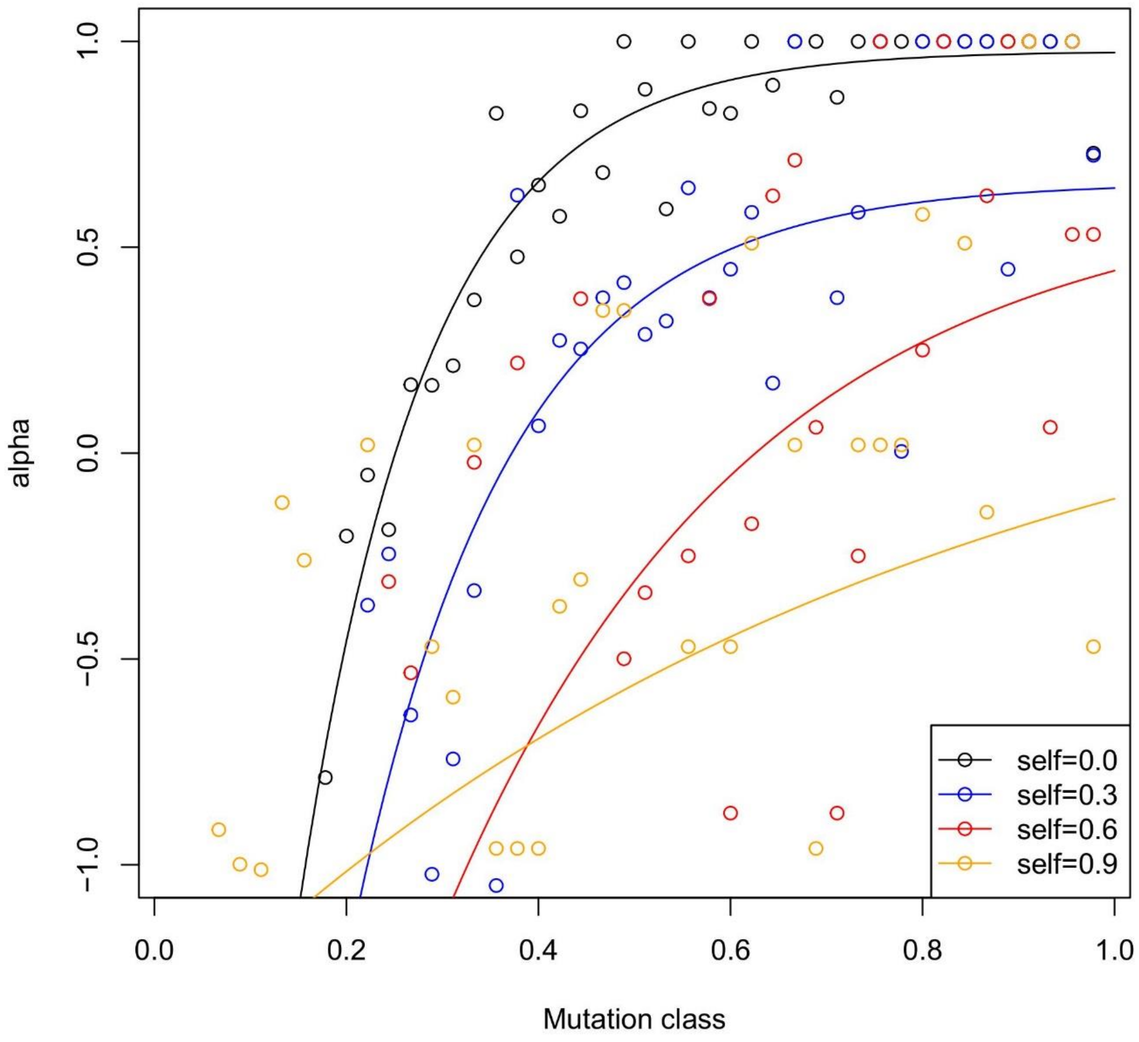


Figure S5: Analysis of the impact of different selfing conditions on the convergence of the asymptote and the overall α estimation. The X-axis indicates different SFS classes and the Y-axis indicates the α estimates. The SFS class-specific α estimates are indicated in hollow coloured circles, different colours are indicative of different selfing intensities.
